# APOE4 Drives Uniquely Dysfunctional Human Microglial States in Alzheimer’s Disease

**DOI:** 10.64898/2026.06.18.733295

**Authors:** Rachel Ee, Meelad Amouzgar, Jumana Afaghani, Kausalia Vijayaragavan, Bryan J. Cannon, Dunja Mrdjen, Dmitry Tebaykin, Angie Spence, Cathrine Sant, David Aley, Zhongyi Guo, Kamilla Sedov, Faria Zafar, Kathleen S. Montine, Amalia Perna, Geidy E. Serrano, Thomas G. Beach, Michael Angelo, Birgitt Schüle, M. Ryan Corces, Thomas J. Montine, Sean C. Bendall

## Abstract

Variation in APOE, notably the ε4 allele, profoundly shapes risk and severity of late-onset Alzheimer’s disease (AD), yet how it remodels human microglial states remains unresolved. We combine spatially resolved proteomic profiling with single-nuclear multiomic analyses to define microglial organization across APOE3/3 and APOE4/4 genotypes in AD. Quantifying condition-associated variation across the cellular manifold reveals a continuous landscape of microglial states. APOE4/4 shifts cells toward terminal states marked by loss of homeostatic identity, metabolic disruption, and incomplete acquisition of disease-associated programs. We identify an APOE4/4-enriched population in AD that exhibits inflammatory signaling without effective metabolic or phagocytic engagement, localizing to niches of gliosis and senescence, and coupled to chronic stress adaptation programs. Together with evidence that APOE4/4 potentiates the activation threshold of nascent microglia, these findings establish a unified framework for human microglial state change, linking genetic risk to spatial and molecular organization of immune responses in the AD brain.

**Graphical Abstract.:** APOE4/4 in Alzheimer’s disease reshapes microglial fate along continuous trajectories characterized by proteomic, transcriptional, and epigenetic programs consistent with chronic stress adaptation, alongside distinct composite spatial niches comprised of astrocytic gliosis and cellular senescence.

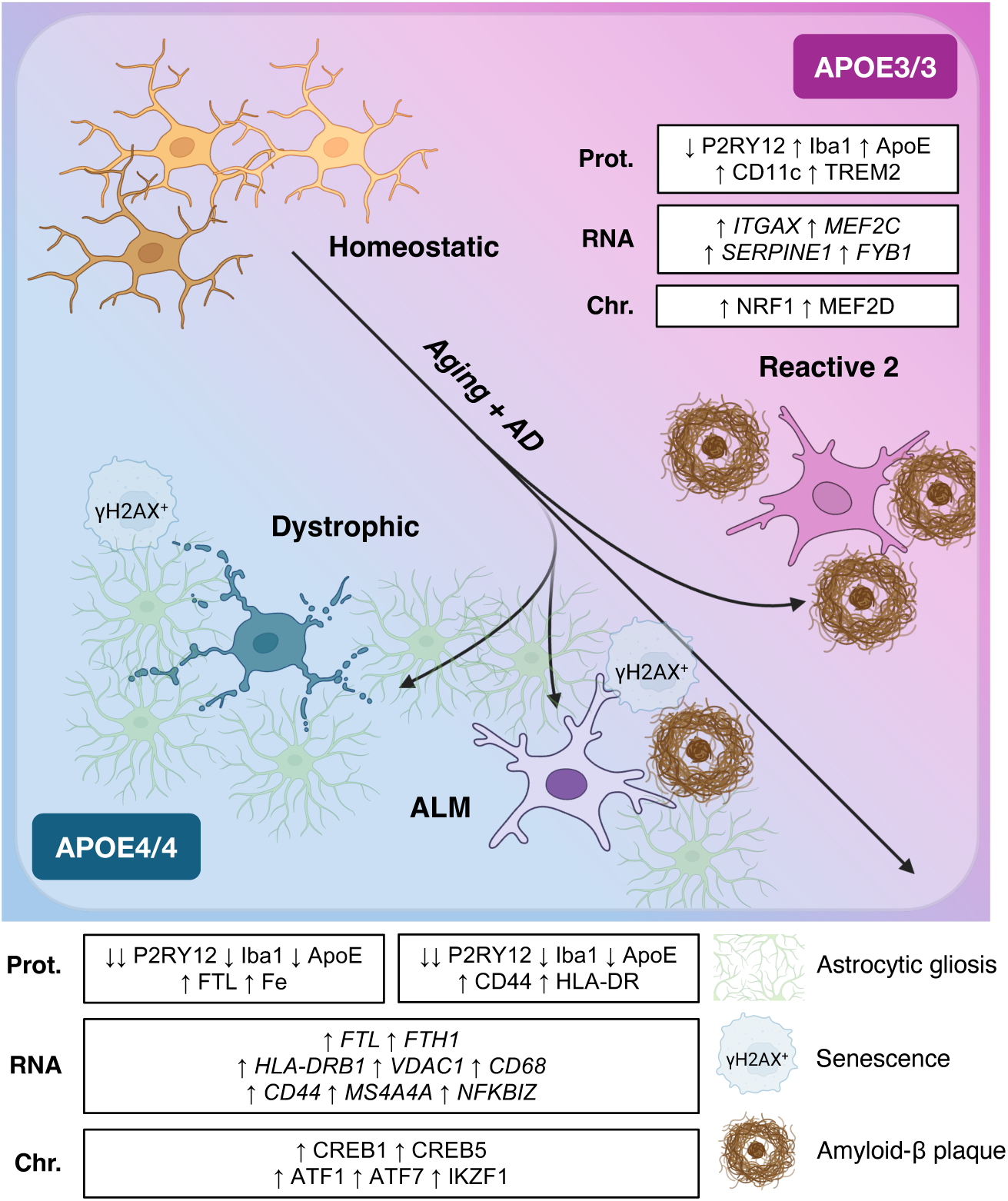

## INTRODUCTION

Alzheimer’s disease (AD) is characterized by the progressive accumulation of amyloid-β (Aβ) plaques and tau pathology (PHF-tau)^1,2^, accompanied by extensive remodeling of the brain’s resident immune compartment: microglia^3^. Genome-wide association studies have established the ε4 allele of apolipoprotein E (APOE4) as the strongest genetic determinant of late-onset AD^4,5^. APOE4, a human variant not native to mouse AD models, exhibits a striking dose-dependent effect, increasing disease risk up to 12-fold in homozygous carriers relative to APOE3 homozygotes^6^ and shifting age of symptom onset earlier in a highly predictable manner^7,8^. Neuropathological studies further demonstrate near-complete penetrance of AD pathology in APOE4 homozygotes across lifespan, exemplified by increased amyloid burden^9^. Beyond its canonical role in lipid transport, APOE is deeply embedded in microglial biology. Converging evidence from mouse models, human induced pluripotent stem cell systems, and human brain tissue indicate that APOE critically shapes immune activation responses in AD and facilitates microglial state transitions^10–12^.

Despite broad agreement that APOE4 reshapes microglial biology, the nature of this remodeling remains incompletely understood in human disease. Several studies report an APOE4-driven pro-inflammatory state without overt functional loss, in which microglia upregulate phagocytosis-associated and inflammatory gene programs^13^, adopt disease-associated microglia (DAM)-like signatures^14^, and acquire a reactive morphology and cytokine profile with increased phagocytic uptake and interferon signaling^15^. In contrast, other studies describe APOE4 microglia that exhibit inflammatory signaling with concurrent functional deficits, including impaired Aβ clearance^16–18^ and defective antigen presentation^19^. Conversely, others report a more globally attenuated phenotype with broad functional impairments, including diminished brain surveillance and process motility^20^, reduced microglial activation and recruitment to Aβ plaques^21^, and impaired responses to acute neurodegeneration, evidenced by decreased phagocytosis of apoptotic neurons and failure to mount canonical disease-associated programs^22–24^.

Resolving these inconsistencies requires a shift in how microglial states are conceptualized. Microglia exist along a continuum of homeostasis, activation, and dysfunction^25^, and are highly responsive to their local microenvironments^26^. However, existing approaches predominantly rely on discrete categorizations of microglial states and analyses of dissociated cells — obscuring both the continuous nature of microglial phenotypes and their spatial context. Moreover, studies examining the effects of APOE variants on microglia often rely on nascent cells derived from pluripotent systems or short-lived animal models, which cannot recapitulate the cumulative influence of decades-long *in vivo* age-associated remodeling that precedes the emergence of clinically apparent AD. As a result, it remains unclear whether the observed heterogeneity reflects intrinsic differences in microglial state, context-dependent responses to local pathology, long-term remodeling that unfolds over decades, or some combination thereof.

Here, we address this gap by integrating high-dimensional spatial proteomics with single-nuclear multiomic characterization of an archival human brain tissue cohort spanning cognitively normal and AD cases on homozygous APOE3/3 or APOE4/4 backgrounds. This approach enabled construction of a spatially resolved map of human microglial states anchored in deep profiling of cellular identity, metabolic remodeling, stress adaptation, and chromatin modifications. Using manifold-based density estimation to quantify condition-associated likelihoods, trajectory inference, and targeted spatial analyses, we define a continuous axis of microglial variation associated with APOE genotype and AD pathology. We identified distinct APOE4-enriched microglial terminal states, including a previously uncharacterized population that we term activation-limited microglia (ALMs), which exhibits features of inflammatory activation yet fail to acquire the full metabolic and phagocytic program characteristic of canonical DAMs. Spatial analysis further demonstrates that these states arise within distinct microenvironmental niches marked by metabolic insufficiency, gliosis, and cellular senescence. Integration with joint single-nuclear transcriptional and chromatin accessibility data revealed that APOE4-associated microglia in AD are characterized by transcriptional and epigenetic programs consistent with chronic stress adaptation, rather than classical acute inflammatory activation. Together, our findings support a model in which APOE4 does not simply induce discrete microglial states, but instead reshapes microglial fate along continuous trajectories, promoting the emergence of maladaptive states defined by incomplete activation and progressive dysfunction. This continuum-based framework reconciles previously conflicting views of APOE4 microglia and provides a unified view of how genetic risk modulates microglial dynamics in AD.

## RESULTS

### Profiling primary human brain tissue across APOE genotype and AD status

Primary human brain tissue cohorts that support direct comparison of homozygous APOE3/3 and APOE4/4 backgrounds across AD status are a scarce resource, especially due to the rarity of cognitively normal APOE4/4 individuals. To profile microglia within their native cellular and pathological landscapes, we leveraged archival formalin-fixed paraffin-embedded (FFPE) autopsy samples from two brain regions: the hippocampal cornu ammonis 1 (CA1) subfield, which is heavily involved in AD neuropathologic change^27–29^, and the cerebellum, which is relatively spared^30,31^. The cohort comprised of 12 cases stratified by APOE genotype and AD status into four groups: APOE4/4 individuals with AD (E4-AD; n = 5, median age = 84.2 years), APOE4/4 cognitively normal individuals (E4-CN; n = 2, median age = 75.5 years), APOE3/3 individuals with AD (E3-AD; n = 3, median age = 78 years), and APOE3/3 cognitively normal individuals (E3-CN; n = 2, median age = 83 years) (**Figure 1A–B; Table S1)**. Inclusion criteria were homozygosity for one of the APOE alleles of interest. Cases had annual clinical assessment of probable AD dementia with high level for AD neuropathologic change, and controls had annual clinical assessment of normal cognition with no or low AD neuropathologic change^32,33^. Exclusion criteria were clinical diagnosis of neurological disease other than AD and any evidence pathologic evidence of neurodegenerative co-morbidity from Lewy body disease, glial cytoplasmic inclusion, pathologic TDP-43 inclusion, hippocampal sclerosis, or any other less common neurodegenerative disease. All cases and controls satisfying these criteria with available tissue blocks were analyzed.

**Figure 1:**
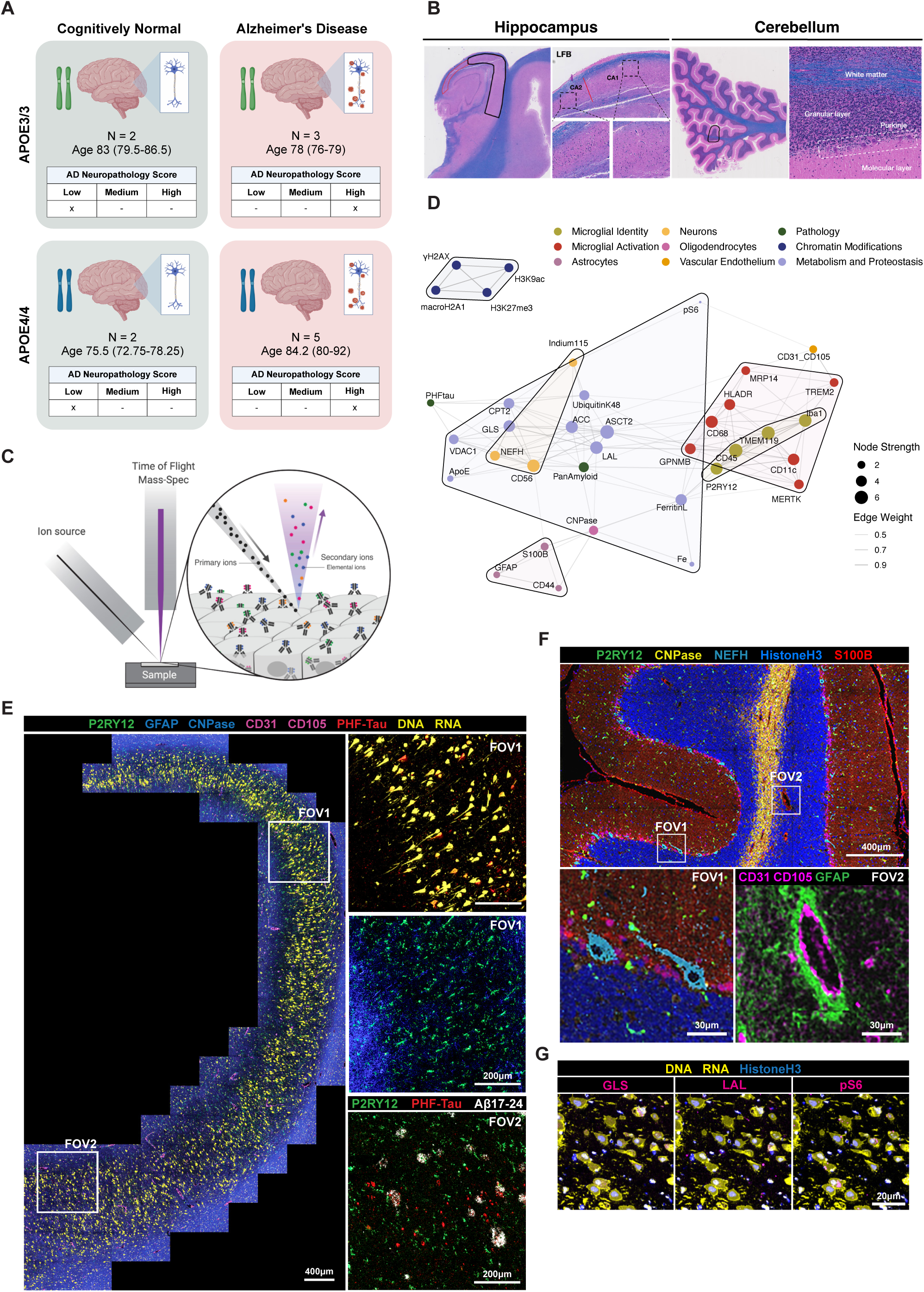
Cohort overview and spatial proteomic mapping of single-cell phenotypes, pathology, metabolism, and histone modifications. **(A)** Cohort overview with *APOE* genotype (rows) and clinico-pathologic diagnosis (columns). Each group tile shows the number of individuals, age at death summarized as median (IQR) in years, as well as distribution across AD neuropathology severity scores. **(B)** MIBI-TOF acquisition workflow includes human brain anatomical region and field-of-view (FOV) selection, staining with metal-conjugated antibody panel, **(C)** image acquisition on the MIBIScope platform, and extraction of 40-dimensional multiplexed images. **(D)** Force-directed network visualization reveals distinct cellular and functional marker constellations across brain tissue. Edges denote significant pairwise co-expression relationships between markers across all segmented objects (Spearman rank correlation coefficient (ρ), FDR ≤ 0.05, |ρ| ≥ 0.3), with edge thickness and opacity proportional to correlation magnitude. Node size reflects node strength (sum of all weighted correlations connected to that marker), indicating its relative centrality within the network. Hulls visually cluster markers into functional constellations, highlighting the variety of biological modules that span the multiplexed imaging panel. **(E)** Representative MIBI image of hippocampus from an E4-AD donor (08-71). An enlarged view of FOV1 highlights fine cellular features including pyramidal neuronal soma and dendrites (yellow), tau-laden neuronal cell bodies (yellow + red), and P2RY12^+^ microglia (green). An enlarged view of FOV2 highlights neuritic plaques that appear as spherical structures consisting of a central core of Amyloid_17-24_ (white), surrounded by neurofibrillary tangles (red). P2RY12^+^ microglia (green) are seen to be proximal to these pathology objects. **(F)** Representative MIBI image of cerebellum from an E3-CN individual (17-07). **(G)** Representative MIBI image demonstrating cytoplasmic localization of select metabolic proteins (GLS, LAL, pS6) within pyramidal neurons. Scale bar denotes 20µm.

In addition to proteomic quantification at single-cell resolution, multiplexed ion beam imaging by time-of-flight (MIBI-TOF) preserves *in situ* spatially resolved interactions and reliably identifies AD parenchymal features^25,34^ (**Figure 1C)**. Our 40-plex antibody panel was designed to capture metabolic, immune, and structural features across major central nervous system cell types (**Figure 1D; Table S2)**. In CA1 gray matter, indium labeling of nucleic acids (yellow) delineated densely packed pyramidal neuronal soma and dendrites forming a compact laminar band (**Figure 1E, left)**. Enlarged fields-of-view (FOVs) highlight pathologic tau-laden neuronal cell bodies (yellow + red) (**Figure 1E, FOV1)** and neuritic plaques composed of a central Aβ core (white) surrounded by pathologic tau-positive neurites (red) (**Figure 1E, FOV2)**. CNPase (yellow) delineated white and grey matter^35,36^, and together with S100B (red) captured tissue architecture and network features often overlooked in non-spatial assays (**Figure 1F)**. Cytoplasmic localization of metabolic enzymes (**Figure 1G)** and nuclear restriction of histone modifications further confirmed marker specificity and subcellular fidelity **(Figure S1A–B)**. As previously described in Mrdjen et al., 2025^25^, microglia were segmented using an intensity-thresholding approach based on composite expression of pan-microglial markers P2RY12, Iba1, and TMEM119. To exclude small transverse process fragments, percentile-based morphometric filters for eccentricity and major-to-minor axis ratio were applied. Optimal thresholds minimized marker variability and preserved biologically meaningful cell morphologies **(Figure S1C)**. Nucleated cells were identified by area-normalized Histone H3 signal intensity **(Figure S1D)**. Segmentation outputs were visually inspected across various FOVs and cases to confirm accurate single-cell delineation.

To broadly assess coordinated relationships across measured proteins, we constructed a global co-expression network encompassing all segmented objects (**Figure 1D)**. The resulting network displayed a modular organization reflecting both cell-type specificity and inter-module coupling. Microglial identity and activation markers formed a dense, highly interconnected module consistent with coordinated expression across homeostatic and reactive states. Astrocytic markers (GFAP, S100B, CD44) and histone modification markers (H3K9ac, H3K27me3, macroH2A1.2, γH2A.X) similarly formed distinct, internally cohesive modules. In contrast, metabolic features exhibited a more distributed topology, maintaining intra-module correlations while also connecting with microglial and astrocytic nodes, positioning bioenergetic regulation as a shared molecular axis bridging immune activation and AD pathology. Together, these results underscore the interconnected molecular architecture of the human hippocampus.

### APOE genotype alters microglial protein programs

To disentangle the independent effects of APOE genotype and AD status from genotype-by-diagnosis (G×D)-influenced molecular programs across CNS cell types in the hippocampal compartment, we applied a donor-aware mixed-effects framework to all single cells with planned marginal-means contrasts. Continuous marker intensities reflect signal quantified within segmented cellular boundaries and may capture both cell-intrinsic protein and associated extracellular or engulfed material. Since this model averages across cellular heterogeneity, it primarily detects large, consistent shifts across cell populations; accordingly, it served as a global screen prior to higher-resolution, state-resolved analyses. Unsurprisingly, PHF-tau was diagnosis-driven across all three cell types (**Figure 2A)**, with no significant G×D interaction, confirming the model’s sensitivity to established AD pathology **(Figure S2A–B)**. Given the phagocytic capacity of glial cells, this signal likely reflects differences in cellular engagement with tau pathology and phagocytic uptake of tau-containing material rather than intrinsic tau production^37–41^. In contrast, total Histone H3 showed no genotype or diagnosis effects, consistent with its role as a structural control (**Figure 2A)**. While several regulatory histone modifications (macroH2A1.2, γH2A.X, H3K9ac, H3K27me3) displayed significant G×D interactions in neurons (**Figure 2A; S2A–B)**, inspection of single-cell distributions revealed that these effects largely reflected broader dispersion in E3-CN neurons negative for these markers rather than uniform genotype-dependent shifts (**Figure 2B)**. We approximated global chromatin accessibility per nucleated cell using the ratio of H3K9ac to H3K27me3, reasoning that higher relative H3K9ac indicates are more transcriptionally permissive chromatin state^42^, whereas higher relative H3K27me3 reflects a more transcriptionally repressive chromatin state^43^. Unsupervised clustering resolved two neuronal populations with estimated global chromatin compaction (H3K9ac:H3K27me3 < 1) and elevated γH2A.X positivity, consistent with senescent-like states (**Figure 2C)**. Senescent 1 was significantly enriched in AD, with no difference in abundance between E3-AD and E4-AD groups **(Figure S2C)**. Relative to the two other clusters, Senescent 1 neurons showed markedly elevated γH2A.X (magenta) and increased phosphorylated ribosomal S6 (pS6) (cyan) (**Figure 2D–E; S2D)**, consistent with persistent DNA damage and a stress-associated phenotype^44–46^. Thus, neuronal G×D interaction signals primarily reflected the emergence of an AD-associated senescent state rather than APOE genotype-specific neuronal programs in this cohort.

**Figure 2:**
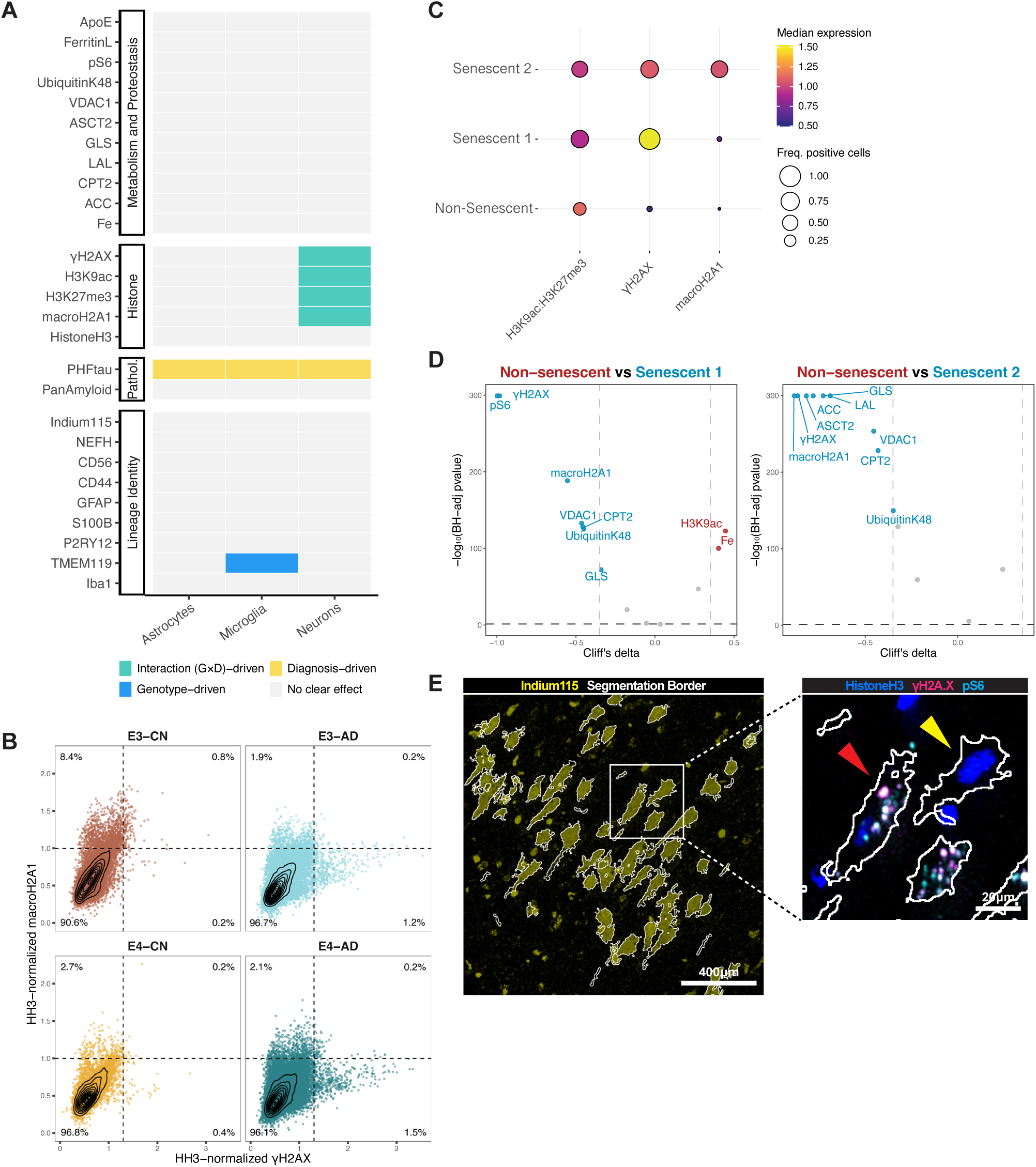
Population-level modeling reveals *APOE*-linked genotype effect confined to microglia. **(A)** Effect classification across markers and cell types. For each marker, a linear mixed-effects model was fit with fixed effects for *APOE* genotype, diagnosis, cell type, and all interactions, with a random intercept for between-individual variability. Outcomes are standardized per marker. Genotype and diagnosis main effects, and the genotype-diagnosis (G×D) interaction were estimated via marginal means with equal weighting across the conditioning factor. Tiles were labelled by an interaction-first hierarchy: interaction-driven when |G×D| ≥ 0.20 SD with FDR < 0.05; otherwise, genotype-driven, or diagnosis-driven when the corresponding main effect was ≥ 0.20 SD with FDR < 0.05 and G×D was not substantive (|estimate| < 0.20 SD or FDR ≥ 0.05). Multiple testing was controlled within effect families (Benjamini-Hochberg, α = 0.05). For pathology-associated markers, signal detected in non-neuronal compartments is interpreted as reflecting cellular association with, or exposure to, extracellular aggregates (including phagocytic uptake or spatial proximity), rather than *de novo* protein expression. **(B)** Nucleated hippocampal neurons from both E3-AD and E4-AD groups harbored a distinct population highly enriched in the DNA-SCARS marker γH2A.X. **(C)** Clustering revealed two senescent subpopulations, termed Senescent 1 and Senescent 2, both of which exhibited global chromatin compaction (H3K9ac:H3K27me3 < 1). **(D)** Senescent 1 was found to show elevated levels of γH2A.X and macroH2A1.2 positivity relative to non-senescent neurons. Senescent 1 also exhibits elevated mTOR signaling (elevated pS6), GLS, and CPT2, consistent with prior reports of metabolic hyperactivity in senescent cells. **(E)** Representative MIBI image of Senescent 1 (red arrow) and non-senescent (yellow arrow) neuron from E4-AD hippocampal sample (11-107). Scale bar on left denotes 100µm; scale bar on right denotes 20µm.

In contrast, an APOE genotype-driven effect was observed in microglia: TMEM119 expression was significantly reduced in APOE4 compared with APOE3 individuals (β_(E4-E3)_ = –0.717, 95% CI = [–1.15, – 0.289]; FDR = 0.028), independent of diagnosis and without a G×D interaction (**Figure 2A; S2A–B)**. Within the scope of this study, population-level APOE effects were most readily observed in microglia. These findings motivated subsequent granular single-cell analyses of microglial state dynamics to dissect APOE-dependent influence in AD.

### Microglial states span a continuum of activation and dystrophy

To quantify the effect of APOE genotype within the microglial single-cell proteomic manifold across CN and AD cases, we utilized MELD^47^ (Manifold Enhancement of Latent Dimensions), a graph signal processing approach that infers the relative likelihood of observing a given cell in one of four APOE-diagnosis conditions. These condition-associated likelihoods – henceforth denoted *p*(condition) – revealed continuous likelihood gradients across the manifold, including regions with elevated *p*(E4-AD) suggestive of distinct cell states underlying the microglial response to APOE4-associated neurodegeneration (**Figure 3A)**.

**Figure 3:**
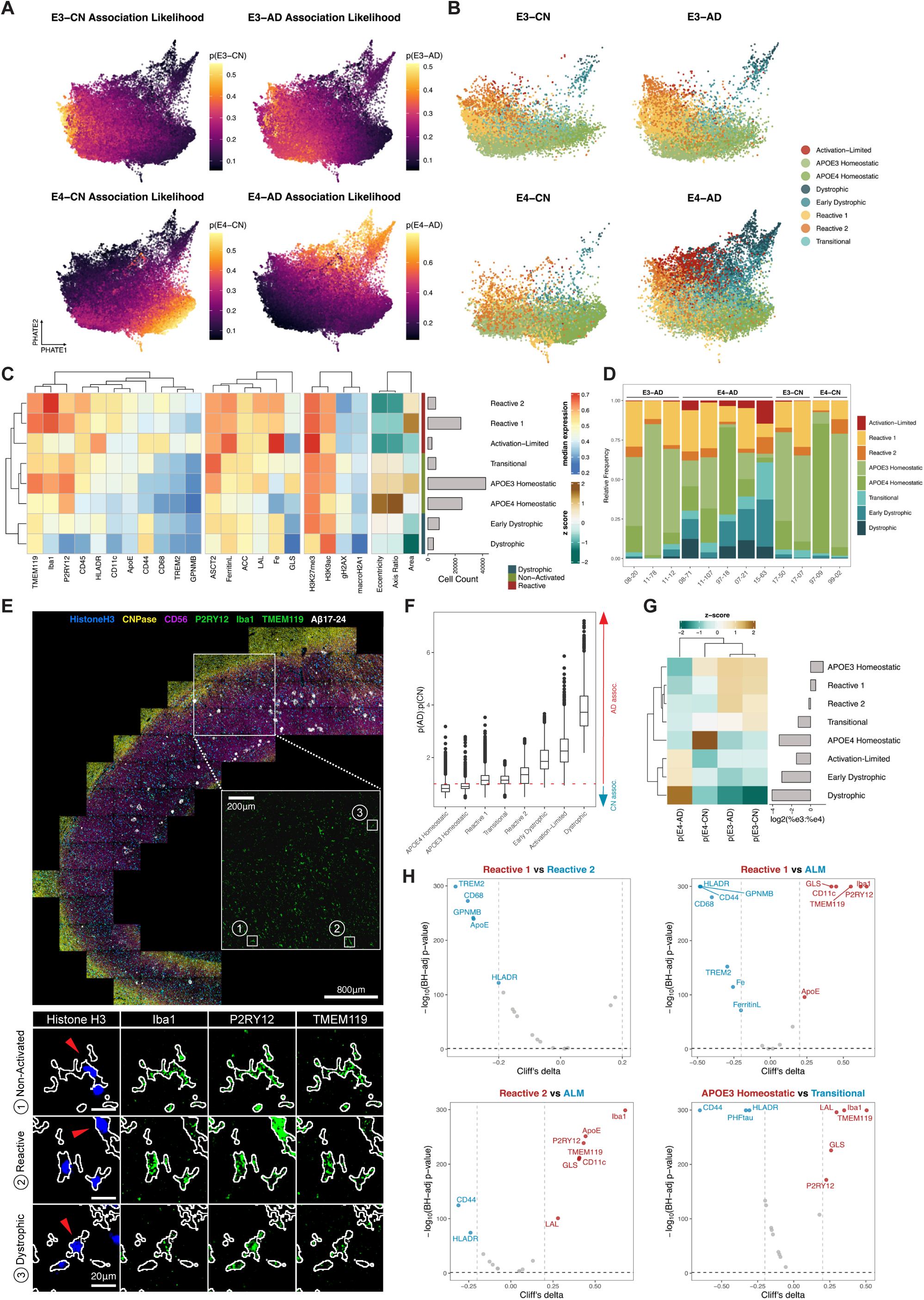
APOE genotype and AD diagnosis jointly shape microglial state composition. **(A)** PHATE embedding of hippocampal microglia colored by MELD-derived condition-association likelihood and faceted by APOE-diagnosis group. High E4-AD likelihood concentrates within a distinct region of the latent space, marking cell states strongly associated with E4-AD. **(B)** Gaussian Mixture Model-based soft clustering resolves eight microglial clusters. **(C)** Expression heatmap of key microglial markers (TMEM119, P2RY12, CD68, L-Ferritin) and morphological parameters (eccentricity, area, axis ratio) used to classify clusters into biologically interpretable states. **(D)** Relative abundances of all clusters per individual, stratified by APOE-diagnosis group. **(E)** Representative multiplexed images of microglia from each phenotype group. Single-cell microglial objects were segmented using an intensity thresholding-based approach on a composite channel consisting of pan-microglial markers (P2RY12, Iba1, TMEM119). Scale bars in multi-panel single-cell callout images (bottom) represent 20µm. **(F)** Cluster-level *p*(AD):*p*(CN) distributions. Both homeostatic states have interquartile ranges below 1, consistent with preferential association to cognitively normal (CN) individuals. The red dotted line indicates *p*(AD):*p*(CN) = 1. **(G)** Cluster-level MELD likelihood estimates (z-scored, left) and log_2_ fold-change in prevalence between APOE3/3 and APOE4/4 individuals (right), highlighting expansion of Transitional, Activation-Limited, Early Dystrophic, and Dystrophic states in APOE4/4. **(H)** Volcano plots showing differential marker expression between clusters.

To annotate the manifold space with biologically interpretable labels, we applied unsupervised clustering using a Gaussian Mixture Model (GMM) framework and identified eight microglial populations: APOE3 Homeostatic, APOE4 Homeostatic, Reactive 1, Reactive 2, Transitional, Activation-Limited, Early Dystrophic and Dystrophic (**Figure 3B–C)**. Stratifying cluster abundances by APOE-diagnosis group revealed increased representation of the latter four across E4-AD individuals (**Figure 3D)**. The three largest populations by cell count were APOE3 Homeostatic, APOE4 Homeostatic and Reactive 1 (**Figure 3C)**. Based on pan-microglial (TMEM119, P2RY12) and activation indicators (CD68, L-Ferritin), clusters were grouped into three higher-order phenotypic categories: <u>Non-Activated</u> (APOE3 Homeostatic, APOE4 Homeostatic, Transitional), <u>Reactive</u> (Reactive 1, Reactive 2, Activation-Limited), and <u>Dystrophic</u> (Early Dystrophic, Dystrophic). TMEM119 and P2RY12 defined the homeostatic axis, with TMEM119^high^ P2RY12^high^ cells representing the non-activated population^48–50^ **(Figure S3A, left)**. In parallel, CD68 and L-Ferritin expression defined the activation and dystrophic axes. Of the non-homeostatic fraction, cells that were CD68^-^ L-Ferritin^+^ were considered dystrophic^51,52^, while cells that were CD68^+^ L-Ferritin^+^ were considered reactive^53,54^ **(Figure S3A, right)**. Cluster morphology supported these classifications (**Figure 3C)**. Cells with reduced eccentricity and axis ratios, coupled with increased area, resembled ameboid microglia (**Figure 3E, second row)**, consistent with immune activation and loss of ramification^55–57^. Conversely, highly eccentric, elongated cells with intermediate area were reminiscent of surveillant, ramified microglia in a resting state (**Figure 3E, first row)**^55,58^. Finally, cells with both reduced area and reduced morphological complexity were interpreted as dystrophic^59^, reflecting process fragmentation and cellular atrophy (**Figure 3E, third row)**.

Both homeostatic clusters were preferentially associated with cognitively normal status (*p*(CN) > *p*(AD)) (**Figure 3F)**. Within the combined homeostatic population, APOE3 Homeostatic cells were preferentially enriched in APOE3/3 individuals, whereas APOE4 Homeostatic cells were enriched in APOE4/4 individuals **(Figure S3B)**, indicating genotype-dependent redistribution within the homeostatic compartment. The APOE3 Homeostatic population best represented the canonical homeostatic phenotype in this dataset. APOE4 Homeostatic microglia exhibited reduced P2RY12 and H3K9ac levels, consistent with prior reports of altered baseline microglial states in APOE4^24^. Genotype-associated shifts were also evident in non-homeostatic states. Transitional, Activation-Limited, Early Dystrophic, and Dystrophic clusters were more prevalent in APOE4 individuals, whereas Reactive 1 and Reactive 2 showed minimal genotype bias (**Figure 3G, right)**. Consistently, Early Dystrophic, Activation-Limited, and Dystrophic clusters displayed high *p*(E4-AD) likelihoods (**Figure 3G, left)**, identifying them as characteristic APOE4-associated disease states.

Targeted comparisons within phenotypic groups were performed to characterize APOE4-associated expression programs. Amongst non-activated microglia, the Transitional cluster was compared with APOE3 Homeostatic cells to capture molecular changes accompanying deviation from the canonical homeostatic state. Amongst reactive microglia, Activation-Limited cells (high *p*(E4-AD)) were contrasted with genotype-neutral reactive states (Reactive 1/2) to isolate APOE4-associated inflammatory programs. Together, these results reveal a continuum of microglial states spanning homeostasis, activation, and dystrophy, and highlight APOE genotype-dependent differences in the composition of these populations in AD.

### Transitional microglia reflect erosion of homeostatic identity in E4-AD

The Transitional cluster was preferentially enriched in the E4-AD group (**Figure 3D; S3C)**, despite its lack of strong condition-specific likelihood estimates (**Figure 3G)**. Hierarchical clustering of median feature expression profiles placed this cluster as the closest neighbor to the APOE3 Homeostatic population (**Figure 3C)**, suggesting a phenotypically intermediate identity. However, compared to APOE3 Homeostatic microglia, the Transitional population exhibited reduced expression of core microglial identity features (Iba1, TMEM119, and P2RY12) alongside elevated expression of HLA-DR, CD44, and PHF-tau (**Figure 3H, bottom right)**. These observations suggest that the Transitional phenotype represents an APOE4-enriched state marked by partial erosion of homeostatic surveillance capacity^48,49^ and an incipient shift toward antigen presentation and early activation^53,60,61^ potentially coupled to tau-associated stress.

### Activation-Limited Microglia (ALMs) represent an E4-AD-enriched reactive state distinct from DAMs

Relative to the APOE3 Homeostatic population, Reactive 1 microglia showed increased expression of microglial activation features TREM2 and CD11c, consistent with an early reactive, actively surveillant phenotype^62^ (**Figure 3C; S3D)**. Reactive 2 microglia demonstrated further activation, with elevated expression of TREM2, CD68, GPNMB, HLA-DR and ApoE, together with marked reductions in homeostatic markers P2RY12 and TMEM119 – reminiscent of late-stage disease-associated microglia (DAMs)^11,63^ (**Figure 3H, top left)**. Notably, neither Reactive 1 nor Reactive 2 differed in abundance between the APOE-diagnosis groups **(Figure S3C)**. Thus, while late-stage DAMs are associated with AD (**Figure 3F)**, they do not account for the increased disease burden observed in APOE4 homozygotes.

Conversely, the Activation-Limited cluster – hereafter referred to as Activation-Limited Microglia (ALMs) – was differentially enriched in E4-AD relative to E3-AD (**Figure 3D; 3G; S3C)**. ALMs exhibited elevated expression of several activation markers, including TREM2, L-Ferritin, CD68, HLA-DR, CD44 and GPNMB, relative to APOE3 Homeostatic and Reactive 1 (**Figure 3H, top right, S3E)** microglia. Despite this heightened immune activation profile, ALMs were molecularly distinct from the DAM-like Reactive 2 population. Specifically, ALMs showed stronger loss of homeostatic identity (P2RY12, TMEM119) while failing to engage the lipid-processing and phagocytic programs characteristic of canonical DAMs^10,11^, reflected by diminished ApoE, LAL and CD11c expression (**Figure 3H, bottom left)**. Notably, ALMs displayed upregulation of HLA-DR (**Figure 3H)**, concordant with prior single-nucleus RNA sequencing studies reporting elevated *HLA-DRB1* expression in APOE4/4 relative to APOE3/3 AD microglia^18^. Together, these findings suggest that ALMs represent a reactive state in which inflammatory activation occurs without the full metabolic and phagocytic reprogramming that typically accompany the canonical DAM transition.

### Distinct pathological niches define microglial states

To dissect how the local tissue environment influences microglial state heterogeneity, we quantified pixel- and object-level spatial features within a 20-pixel (7.8µm) radius around each microglial cell. Pixel-based spatial features were assigned using feature-intensity thresholds derived from microenvironment-level expression profiles, including Fe for free iron, CD44/GFAP co-expression for astrocytic gliosis, and CNPase for myelin-rich regions. In parallel, object-based spatial features were defined by the presence of segmented object(s) within a given microenvironment, enabling proximity-based associations between individual microglia and nearby Aβ plaques, PHF-tau⁺ neighbors, and γH2A.X⁺ nucleated cells, used here as a proxy for senescence^64,65^. Together, these features captured both local expression of metabolic features and proximity to pathological cues including astrocytic gliosis, tauopathy, Aβ plaques, free iron, and cellular senescence – thus generating a unique microenvironmental profile for each cluster. Across clusters, microglial states occupied distinct pathological niches (**Figure 4A)**. Homeostatic microglia exhibited minimal associations with pathologic spatial features and the lowest median neighborhood Shannon entropy, aligning with their expected residence in relatively undisturbed tissue with low local diversity, rather than pathology-enriched microenvironments (**Figure 4A–B; S4A)**. Consistent with their DAM-like activation profile, Reactive 2 microglia were most strongly enriched for Aβ plaque association (purple), followed by ALMs (**Figure 4A–C; S4B)**. Comparison of plaque-associated cells between these two states revealed that ALMs resided at significantly greater distances from plaques compared to Reactive 2 microglia **(Figure S4C)**, suggesting ALMs are less directly responsive to proximal Aβ and reside in the broader plaque-adjacent environment. Instead, ALMs were more frequently localized to GFAP⁺CD44⁺ gliosis zones (green), and regions containing γH2A.X⁺ (blue) and PHF-tau⁺ (red) neighbors (**Figure 4A–C; S4A; S4D)**, indicating a distinct inflammatory niche composed of non-Aβ stressors. Microglial recruitment to plaques was quantified as an enrichment ratio (log_2_ observed/expected microglial area) within each plaque’s 50-pixel expansion zone relative to a uniform distribution across the FOV. Overall, microglia in E3-AD were more spatially concentrated around plaques than in E4-AD **(Figure S4E)**, further reinforcing the fact that APOE4 alters not just cell-intrinsic microglial states, but their spatial targeting of pathology.

**Figure 4:**
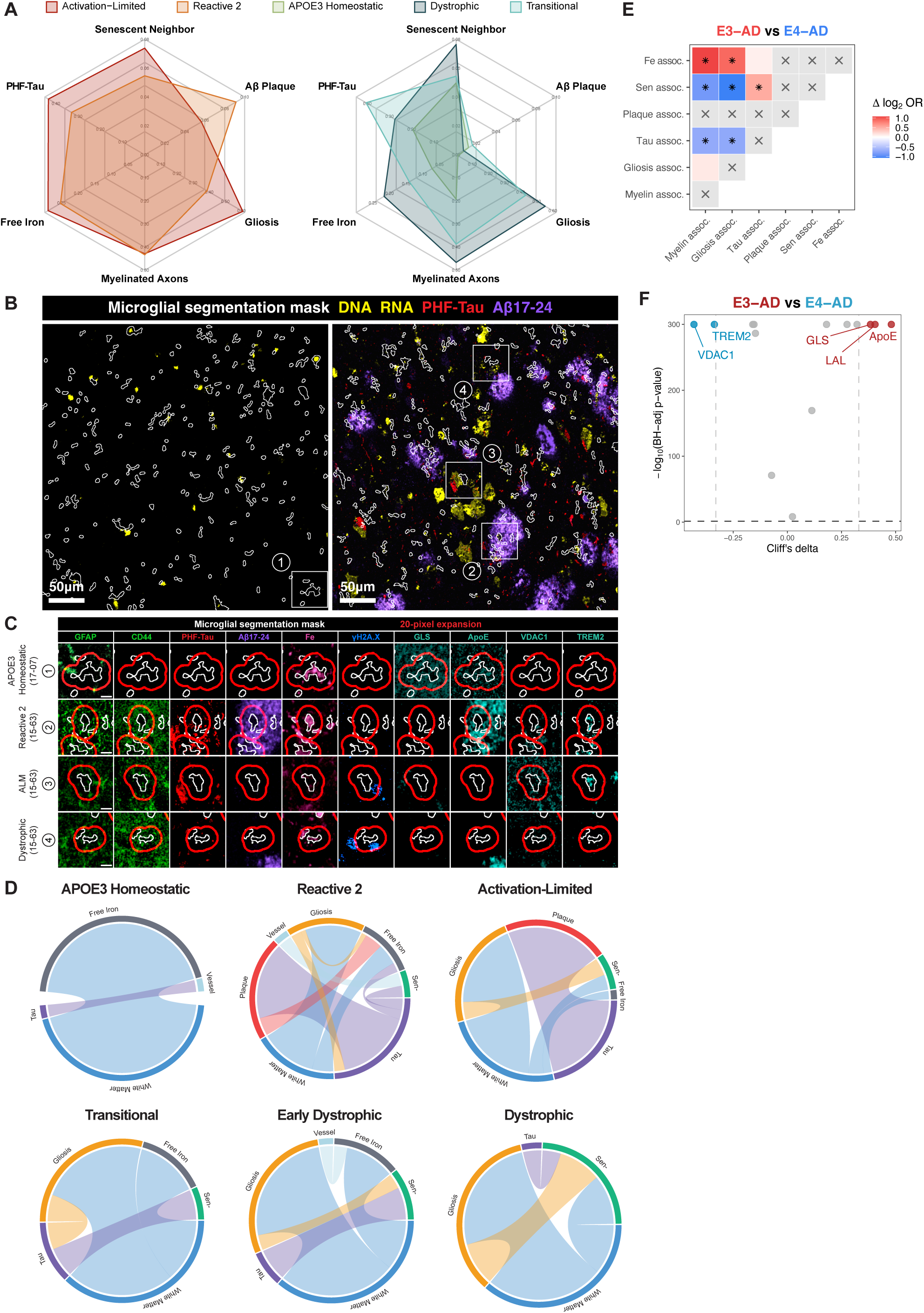
E4-AD microglial niches are defined by hypometabolism and senescence-gliosis co-occurrence. **(A)** Proportions of microglia associated with key spatial features across clusters, including myelinated axons, PHF-tau^+^ neighbors, amyloid plaques, astrocytic gliosis, free iron, and senescence. Astrocytic gliosis was assessed via mean pixel expression of GFAP and CD44 within each cell’s 20-pixel expansion region; white matter and free iron association was similarly defined using CNPase and Fe expression respectively. **(B)** Left: Multiplexed image of a CA1 field-of-view from an E3-CN individual (17-07), highlighting the absence of overt tau and amyloid pathology. A microglial cell labeled 1 corresponds to the homeostatic microglia shown in (D). Right: Multiplexed image of a CA1 field of view from an E4-AD individual (15-63), showing extensive accumulation of compact amyloid plaques (purple) and PHF-tau accumulation (red). Microglial cells labeled 2–4 correspond to the plaque-associated Reactive 2, senescence-associated ALM, and Dystrophic microglia shown in (D). Scale bars represent 50µm. **(C)** Representative multiplexed images highlighting ALM and Dystrophic microglia embedded in astrocytic scarring, with proximal senescent neighbors. Dense CD44^+^GFAP^+^ meshwork (green) and γH2AX^+^ cells (blue) highlight the convergent pathology shaping the local microglial niche. Scale bars represent 10µm. **(D)** Cluster-specific co-occurrence networks of spatial features. Edges represent significant, positively enriched feature pairs (FDR < 0.05), with edge thickness proportional to log_2_ odds ratio (OR). Low-prevalence features (<5% per cluster) were excluded to improve interpretability. Pairwise associations were tested using Fisher’s exact test on 2×2 contingency tables, with multiple testing corrections applied within each cluster. **(E)** Heatmap of differential co-occurrence between E3-AD and E4-AD groups, showing Δlog₂OR for each spatial feature pair. A 2×2×2 log-linear model tested whether the strength of association between each feature pair differed by genotype. Stars denote FDR-adjusted significance. **(F)** Comparison of metabolic environment between E3-AD and E4-AD microglia.

To better understand how pathological features organize across the hippocampal landscape, we examined their global pairwise co-occurrence relationships **(Figure S4F)**. As internal controls, Aβ plaques and myelinated regions showed strong mutual exclusivity while pathologic tau and Aβ plaques showed robust co-occurrence as expected. We next examined pairwise feature co-occurrence within the local microenvironments of each cluster (**Figure 4D)**. Reactive 2 microglia exhibited widespread feature interactions, including (Aβ, PHF-tau), (gliosis, myelinated axons), and (senescence, vessel) – consistent with Aβ plaque-centric inflammatory signaling^66–68^ and vascular interaction in advanced AD^69,70^. In contrast, ALMs displayed co-occurrence patterns not seen in DAM-like microglial niches, such as (gliosis, senescence) and (myelinated axons, senescence), suggesting a unique convergence of multiple pathological stressors in their local environment **(Figure S4G)**.

Finally, genotype-stratified analysis revealed divergent spatial networks between APOE genotypes in AD (**Figure 4E)**. E4-AD microglial niches showed enrichment for co-occurrences linking gliosis, tau pathology, white matter degeneration, and cellular senescence. In contrast, E3-AD microglial niches showed enrichment of (free iron, gliosis) and (free iron, myelinated axons) interactions, indicating divergence in pathological network architecture.

### E4-AD microglia reside in hypometabolic microenvironments

While the above analyses identified the pathological contexts associated with each microglial state, we next asked whether broader metabolic properties of the local microenvironment also contribute to microglial state specification. To address this, we trained a random forest classifier using both object-and pixel-level features to identify global spatial predictors of microglial cluster identity. Pixel-level metabolic features emerged as the strongest predictors of cluster identity **(Figure S4H)**. Top-ranked features included regulatory features of glutaminolysis (GLS), lipid metabolism (LAL, ACC), mitochondrial stress (VDAC1), and inflammatory lipid sensing (ApoE, TREM2), indicating local metabolic tone as a key determinant of microglial state. Genotype-stratified comparison of neighborhood metabolic expression revealed clear APOE-dependent differences. E4-AD microglial niches were enriched in VDAC1 and TREM2 but depleted in key metabolic and lipid-processing markers (GLS, LAL, and APOE) (**Figure 4F)**, indicating that APOE4 microglia are embedded within metabolically compromised local environments likely linked to their unique dysfunction in AD^14^. Overall, these findings highlight that E4-AD microglial niches are characterized by convergence of gliosis and cellular senescence, coupled with impaired lipid catabolism and efflux^71–74^, blunted glutamine utilization^75–78^, and elevated mitochondrial stress^79–81^ – conditions that preferentially support activation-limited and dystrophic microglial states.

### Microglia diverge along distinct trajectories toward APOE4-associated terminal states in AD

Analysis of donor-level cluster frequencies demonstrated a marked depletion of APOE3 Homeostatic microglia and significant enrichment of Transitional, Early Dystrophic, and Dystrophic population in E4-AD relative to E3-AD, indicating a genotype-dependent shift in microglial state composition **(Figure S3C)**. Notably, abundance of the Transitional population was strongly and inversely correlated with that of the APOE3 Homeostatic population (**Figure 5A)**, suggesting a phenotypic continuum in which erosion of surveillance capacity in E4-AD coincides with the emergence of APOE4-associated dystrophy. In congruence with this notion, mixed-effects ordinal trend modeling across the ordered states (APOE3 Homeostatic → Transitional → Early Dystrophic → Dystrophic) revealed a monotonic decline in core microglial markers (P2RY12, TMEM119, Iba1), accompanied by increased global chromatin accessibility (H3K9ac:H3K27me3), elevated CD44 expression, and a higher proportion of L-ferritin^+^ cells, a feature associated with microglial dystrophy (**Figure 5B–C)**^53,82,83^. Distribution of condition-associated likelihoods across E3-AD and E4-AD similarly revealed a continuous gradient across these states **(Figure S5A)**.

**Figure 5:**
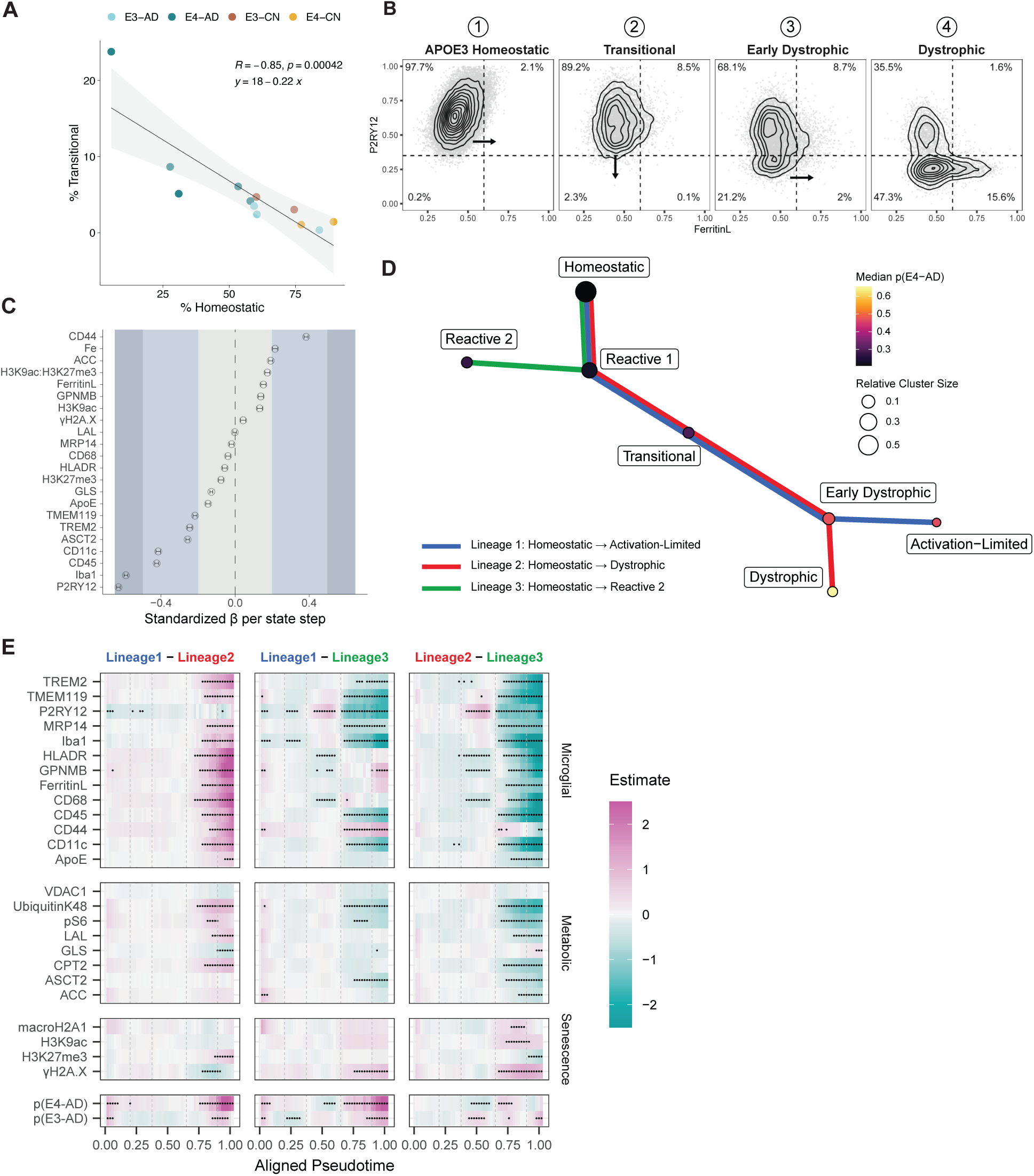
E4-AD-associated trajectories show unique feature expression dynamics across pseudotime. **(A)** Individual-level correlation between Transitional and APOE3 Homeostatic frequencies (R = −0.85, p < 0.001). Shaded gray area denotes 95% CI. **(B)** Biaxial scatter plots of P2RY12 vs L-ferritin, faceted by cluster. The dystrophic state is characterized by a higher prevalence of P2RY12-low, L-ferritin-high microglia. **(C)** Forest plot showing estimated beta coefficients and 95% CI from mixed-effects ordinal trend modeling along the APOE3 Homeostatic → Dystrophic axis. Each row corresponds to a feature tested across the ordered microglial clusters, representing progression from homeostasis toward dystrophy. Positive coefficients indicate that the feature increases stepwise across successive clusters, whereas negative coefficients indicate that the feature decreases stepwise. Case ID was modeled as a random effect to account for inter-individual variability. The vertical dashed line at zero represents the null effect. Shaded regions denote ranges for small, medium, and large effect sizes. **(D)** Graph representation of cluster relationships based on Slingshot-inferred lineage structure and Euclidean distances in lineage weight space. Clusters (nodes) are positioned using a force-directed layout and connected by edges representing the minimum spanning tree (MST) inferred by Slingshot. Node fill indicates the median likelihood estimate of E4-associated AD per cluster. Edge colors represent lineage identity. **(E)** Differential abundance analysis comparing marker expression across lineages within manually defined pseudotime bins. Dots indicate features with significant differences (BH-adjusted p ≤ 0.1).

Together, these findings support a model in which APOE4-associated microglial phenotypes may not represent discrete populations but rather progression along a shared activation continuum. To capture the gradual transitions across metabolic, inflammatory, and epigenetic axes, we applied Slingshot trajectory inference^84^ to order single-cell microglial states in reduced-dimensional space. A graph-based representation of cluster relationships demonstrated this global organization and revealed three partially overlapping lineages emerging from a common root, thus providing a natural starting point for trajectory interpretation (Figure 5D). From this shared origin, Lineages 1 and 2 progressed toward distinct E4-AD-enriched terminal states – ALMs and Dystrophic microglia, respectively – whereas Lineage 3 terminates in the DAM-like Reactive 2 state. While ALMs, Early Dystrophic, and Dystrophic cell states emerged predominantly at late pseudotime, the early and mid-pseudotime regions were largely defined by the Homeostatic population and ensuing emergence of Transitional microglia **(Figure S5B)**. This pattern suggests that lineage divergence originates within broadly shared starting states and becomes apparent only as cells progress along pseudotime.

To comprehensively characterize lineage-specific programs underlying these transitions, we predicted feature expression along aligned pseudotime using generalized additive modeling **(Figure S5C)** and quantified inter-lineage divergence by cosine dissimilarly between aligned expression profiles **(Figure S5D)**. This revealed a shared early state program across all three lineages, with all three lineages subsequently adopting distinct terminal profiles **(Figure S5E)**. Overlaying MELD-derived likelihoods onto the trajectory structure further showed that E4-AD association increased toward late pseudotime across all lineages, with the strongest enrichment in Lineage 1 (ALM) terminal states **(Figure S5F)**.

Differential abundance testing across pseudotime bins identified Lineage 3 (Reactive 2) as the earliest diverging branch at mid-pseudotime, indicating that induction of inflammatory features (HLA-DR, GPNMB, CD68) and reduced P2RY12 expression marks the first major transition away from the shared microglial trajectory (**Figure 5E, middle and right)**. Following this initial divergence, Lineages 1 (ALM) and 2 (Dystrophic) progressively lost canonical microglial identity markers (P2RY12, TMEM119, Iba1), along with diminished expression of TREM2 and CD11c (**Figure 5E, middle and right)**. These changes were accompanied by a decline in metabolic and proteostatic capacity, evidenced by reduced K48-linked ubiquitination, lysosomal acid lipase (LAL), and glutamine transporter (ASCT2) expression. In parallel, indicators of persistent genomic stress and senescence, γH2A.X^64,65^ and macroH2A1.2^85^, increased. Despite this shared loss of homeostatic identity, the two trajectories diverged in their inflammatory programs. Lineage 1 (ALM) retained inflammatory and iron-handling markers (HLA-DR, GPNMB, L-ferritin, CD68, ApoE), in contrast to Lineage 2 (Dystrophic), which saw rapid downregulation of these features beyond mid-pseudotime (**Figure 5E, left; S5G)**.

Collectively, these lineage-specific programs indicate that E4-AD biases microglial trajectories toward terminal states characterized by progressive loss of homeostatic maintenance and incomplete damage-response programs, consistent with cellular exhaustion or dysfunction.

### Orthogonal multiomics recapitulates APOE-associated microglial continuum

To independently validate the APOE-associated microglial continuum observed in our spatial proteomic and trajectory analyses, we analyzed an orthogonal single-nuclear multiome dataset consisting of matched snRNA-seq and snATAC-seq profiles from hippocampal microglia isolated from eight AD donors (n = 4 APOE3/3; n = 4 APOE4/4) **(Table S3)**. RNA and chromatin accessibility profiles were integrated into a unified MultiVI framework^86^, which learns a shared latent representation of cellular state across both modalities. Integration effectively removed donor-driven variation while preserving genotype-associated structure in the joint RNA-ATAC latent space **(Figure S6A)**. Leiden clustering identified minor contaminating cell types. Clusters corresponding to highly proliferative cells and T cells **(Figure S6B)** were excluded to restrict downstream analyses to resident microglia.

To quantify genotype enrichment across the integrated manifold, we applied MELD^47^ to the MultiVI neighborhood graph and derived a genotype difference score representing a continuous measure of APOE4 versus APOE3 association (MELD_Δ_ = MELD_4/4_ - MELD_3/3_). When projected onto the integrated embedding, MELD_Δ_ formed a likelihood gradient across the microglial manifold rather than defining a discrete APOE-specific cluster (Figure 6A), mirroring the continuous activation trajectory inferred from our proteomic lineage analysis. Partitioning MELD_Δ_ values using a GMM framework (k = 3) identified MELD-low, intermediate, and MELD-high populations. While MELD-high cells were enriched for APOE4/4 individuals and MELD-low cells for APOE3/3 individuals, all cases were represented across multiple bins **(Figure S6C)**. These observations indicate that APOE genotype indeed impacts microglial cellular state in this shared multiomic space.

**Figure 6:**
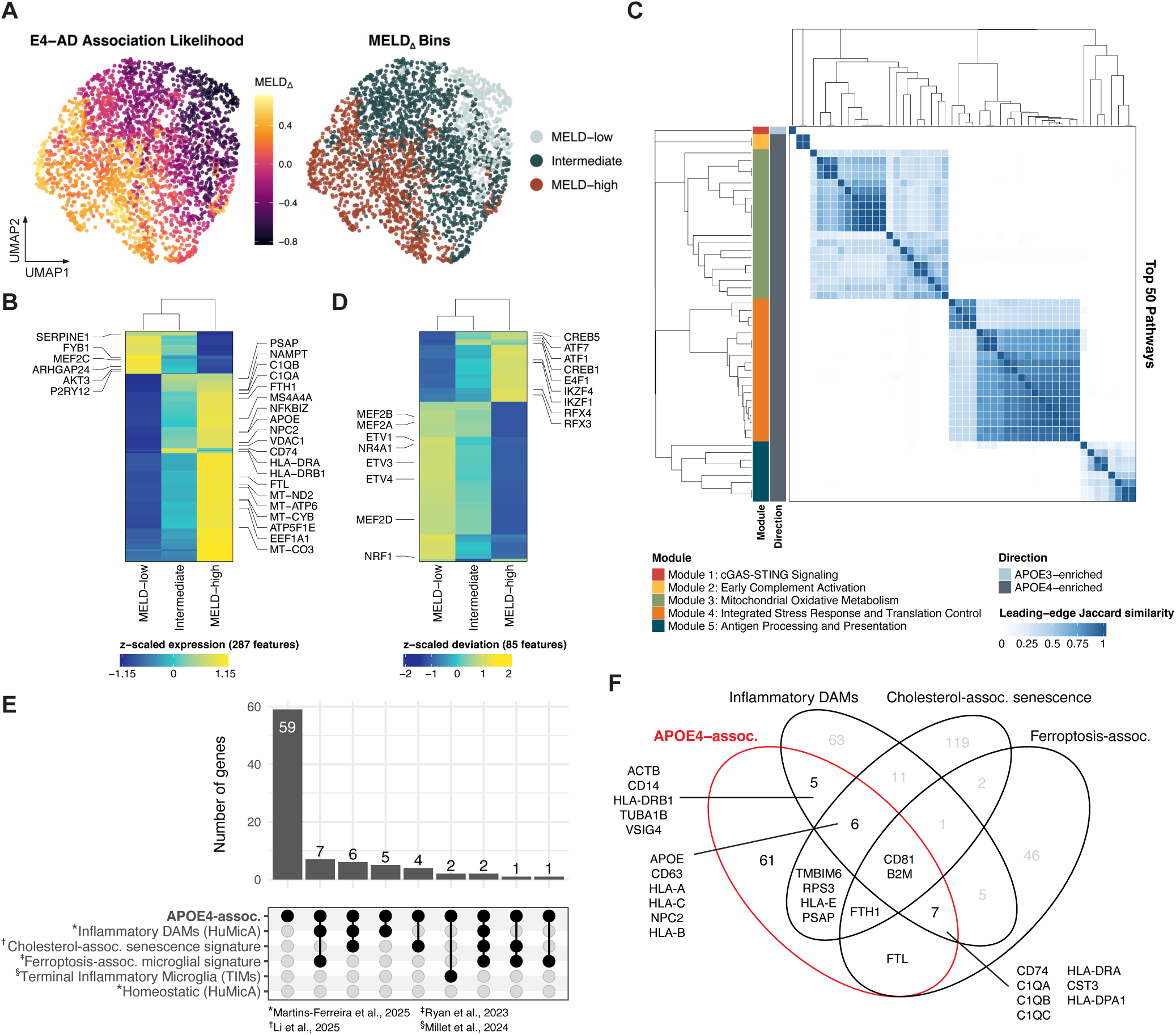
Single-nuclear multiomics reveal a continuous APOE-associated microglial axis with coordinated transcriptional and chromatin remodeling. **(A)** UMAP visualization of microglia embedded in the shared latent space learned by MultiVI from paired RNA and ATAC modalities. Cells are colored by MELD_Δ_ (left) and bin assignment (right), highlighting a continuous APOE-associated gradient across the manifold. **(B)** Heatmap of mean gene expression across MELD_Δ_ bins. Genes shown were identified from donor-aware continuous MELD_Δ_ association analysis (FDR < 0.05). Genes enriched in MELD-low microglia include homeostatic and regulatory factors such as P2RY12, and MEF2C (left). In contrast, MELD-high microglia exhibit increased expression of stress- and activation-associated genes including FTH1/FTL, HLA-DRB1, MS4A4A, and mitochondrial metabolism modules (right). Intermediate bins display graded, intermediate expression patterns, consistent with a continuous transcriptional transition. **(C)** Pairwise similarity between enriched pathways (FDR < 0.05) was computed based on overlap of leading-edge genes derived from GSEA of MELD_Δ_-associated transcriptional changes. The heatmap displays pairwise pathway Jaccard similarity (fill), where higher values indicate greater overlap in leading-edge genes. Both rows and columns represent pathways, ordered by hierarchical clustering of the similarity matrix. The accompanying dendrograms group pathways into higher-order modules, reflecting shared transcriptional programs. Modules were annotated based on the dominant biological processes represented within each cluster. Row annotations indicate module assignment and the direction of enrichment for each pathway. **(D)** Heatmap of mean motif deviation scores across MELD_Δ_ bins for differentially accessible motifs (FDR < 0.05). MELD-low microglia show increased accessibility of motifs associated with homeostatic and regulatory programs (left), whereas MELD-high microglia are enriched for stress-responsive ATF/CREB family motifs (right), consistent with activation of integrated stress response pathways. **(E)** Upset plot showing interactions between genes positively associated with MELD_Δ_ (FDR < 0.05, log2FC ≥ 0.2) and reference microglial transcriptional signatures derived from Martins-Ferreira et al., 2025^103^ (Human Microglia Atlas), Li et al., 2025^104^ (microglial cholesterol-associated senescence signature), Ryan et al., 2023^94^ (microglial ferroptosis-associated signature), and Millet et al., 2024^17^ (Terminal Inflammatory Microglia). Only sets containing overlap with APOE4-associated genes are shown. **(F)** Venn diagrams highlighting overlap between APOE4-associated genes (red), inflammatory DAM signature, microglial cholesterol-associated senescence signature, and microglial ferroptosis-associated signature.

### APOE4-associated microglia exhibit transcriptional programs of chronic cellular stress

Donor-aware pseudobulk differential expression analysis comparing MELD-high and MELD-low populations identified a coordinated APOE4-associated transcriptional signature (**Figure 6B; S6D)**. Consistent with immune activation, APOE4-associated microglia upregulated expression of complement (*C1QA/B/C*) and MHC-II antigen presentation machinery genes (*HLA-DRA*, *HLA-DRB1*, *HLA-DPA1*). However, concurrent upregulation of *MS4A4A*, a negative regulator of TREM2 signaling^87^, together with persistent NF-κB activation marked by *NFKBIZ*^88^, suggested that this inflammatory state may be linked to constraints on canonical TREM2-dependent microglial programs. Additionally, this signature extended beyond canonical immune activation, spanning several functional modules consistent with chronic cellular stress. This included coordinated upregulation of genes involved in mitochondrial oxidative phosphorylation (*MT-ND1/2/3/4*, *MT-CO1/2/3*, *MT-ATP6*, *MT-CYB, ATP5F1E, NAMPT*) alongside lipid trafficking (*APOE*, *NPC2*), iron sequestration (*FTH1*, *FTL*), and redox-associated programs (*VDAC1, GPX*). Given that profiled cells passed stringent mitochondrial read fraction QC thresholds (<5%), this coordinated upregulation reflects a genuine transcriptional shift rather than a technical artifact, and is suggestive of increased energetic demand and compensatory mitochondrial remodeling in response to prolonged metabolic stress^89–91^. Furthermore, the coupling of lipid handling dysregulation, elevated iron storage capacity, and upregulated lipid peroxide scavenging implicates stress pathways associated with ferroptosis vulnerability within APOE4-associated microglia^92–94^.

This binary comparison between MELD-high and MELD-low populations enabled us to identify transcriptional programs most strongly associated with the extremes of the APOE continuum, revealing that APOE4-associated microglia occupy a stress-adapted, rather than a simply inflammatory state. To determine how these programs varied continuously along the full APOE-associated manifold, and to identify broader pathway-level regulation, we modeled gene expression as a continuous function of MELD_Δ_ while controlling for donor identity (**Figure 6B; S6E)**. Genes were ranked by their MELD_Δ_ association statistics and subjected to gene set enrichment analysis (GSEA). Top pathways (FDR < 0.05) were clustered based on leading-edge gene overlap, thus enabling us to distill enriched pathways into biologically coherent programs. This resolved discrete modules with strong internal concordance, separating APOE3- and APOE4-associated transcriptional programs (**Figure 6C)**. A prominent APOE4-enriched module captured the integrated stress response (ISR) and translational control, including GCN2-mediated amino acid starvation signaling, SRP-dependent co-translational protein targeting to the endoplasmic reticulum, and cellular responses to nutrient deprivation (**Figure 6C; S6F)**. This was accompanied by a module characterized by oxidative phosphorylation and neurodegeneration-associated disruptions in complex I function, indicating dysregulated mitochondrial respiration and bioenergetic homeostasis. Together, these findings suggest chronic metabolic and proteostatic stress along the APOE4-associated end of the continuum, in contrast to the innate immune sensing programs (i.e., cGAS-STING signaling) preserved at the APOE3-associated end.

### Stress-responsive regulatory networks underlie the APOE4-associated chromatin landscape

We next asked whether chromatin accessibility differences accompany the transcriptional programs observed along the APOE continuum. Although differential accessibility between MELD-high and MELD-low microglia was modest and involved a relatively small fraction of peaks **(Figure S6G)**, genome-wide chromatin accessibility patterns across transcription factor (TF) motifs revealed coordinated regulatory changes along the APOE continuum. Motif activity analysis using chromVAR^95^ identified enrichment of several TF families in APOE4-associated microglia (Figure 6D**; S6H)**, including CREB/ATF (CREB5, CREB1, ATF1, ATF7), NFAT (NFATC4), and Ikaros zinc-finger family TFs (IKZF1, IKZF4). Enrichment of NFAT and CREB/ATF-family motifs is consistent with persistent calcium- and stress-responsive transcriptional activation^96,97^, while IKZF and RFX family enrichment suggests stabilization of altered immune regulatory states and antigen presentation programs^98^. Notably, APOE4-associated microglia also exhibited enrichment of E4F1 motifs, a regulator of mitochondrial homeostasis and cellular stress checkpoint signaling^99^. In contrast, APOE3-associated microglia preferentially exhibited enrichment of MEF2-family (MEF2A, MEF2B, MEF2D), NRF1, and NR4A1 motifs, consistent with preservation of homeostatic immune regulation^100^, mitochondrial competence^101^, and controlled inflammation^102^. Collectively, these findings support a model in which APOE3-associated microglia retain a metabolically competent and homeostatically regulated chromatin landscape relative to the chronically stress-adapted, APOE4-associated state.

### Multimodal integration reveals stress-adapted, activation-constrained E4-AD microglial states

To further contextualize E4-AD-associated programs observed in this dataset, genes positively associated with MELD_Δ_ (FDR < 0.05, log_2_FC ≥ 0.2) were compared against reference microglial signatures derived from the Human Microglia Atlas^103^ and Terminal Inflammatory Microglia (TIMs)^17^, cholesterol-associated microglial senescence^104^, and ferroptosis-associated microglial states^94^. Overlaps between our APOE4-associated transcriptional signature and published microglial gene signatures were greater than expected by chance, based on one-sided Fisher’s exact tests performed over the set of 36313 genes tested in the continuous MELD_Δ_ analysis **(Figure S6I)**. APOE4-associated genes showed the greatest overlap with the inflammatory DAM signature, with additional convergence on ferroptosis-associated and cholesterol-associated senescence programs, indicating partial alignment with canonical disease-associated activation (**Figure 6E)**. Shared genes between the APOE4-associated signature and DAM/ferroptosis-associated programs included antigen presentation and complement-related genes such as *CD74*, *C1QA*-*C*, *HLA-DRA*, *HLA-DPA1*, and *CST3* (**Figure 6F)**. In contrast, overlap with the DAM/cholesterol-associated senescence signature was characterized by genes linked to lipid handling and lysosomal biology, including *APOE*, *CD63*, *NPC2*, and MHC-I genes (*HLA-A*, *HLA-B*, *HLA-C*). Notably, the strongest APOE4-associated signals were preferentially shared with ferroptosis- and senescence-related programs rather than the DAM signature alone, including *FTH1* and *FTL* **(Figure S6J)**. Amongst genes overlapping specifically with the senescence-associated signature (**Figure 6F)**, *PSAP* stands out as a lysosomal regulator of glycosphingolipid metabolism^105,106^, thereby pointing to engagement of lipid-processing pathways central to APOE biology. Together, the enrichment of ferritin genes (*FTH1*, *FTL*) alongside lysosomal and stress-associated factors such as *PSAP* and *TMBIM6* suggests that the APOE4-associated state extends beyond a canonical DAM-like program and instead incorporates features of iron dysregulation, lipid-lysosomal stress, and senescence-associated remodeling.

The joint snRNA-ATAC analysis independently recapitulated key features of the microglial trajectories identified in our spatial proteomic dataset. Across both modalities, APOE4-associated microglia partially engage aspects of inflammatory activation but were more strongly defined by stress-adapted programs. These molecular features paralleled the ALM and dystrophic states identified at the protein level, providing orthogonal multimodal evidence that APOE4 shifts microglia along a shared continuum toward an activation-constrained phenotype shaped by chronic stress and altered metabolic regulation, rather than full convergence on previously defined inflammatory microglial signatures.

### iPSC-derived APOE4 microglia exhibit an augmented baseline and heightened reactivity

Our analyses of post-mortem human brain tissue provide insight into microglial states within their native spatial context but are inherently limited to a static snapshot of end-stage disease and advanced age. We therefore sought to determine how APOE shapes microglial activity and responsiveness to stimuli in a controlled early-life context. APOE3 and APOE4 differ by a single amino acid at position 112, with Cys112 in APOE3 and Arg112 in APOE4, a substitution that alters protein domain structure and lipid binding function^107^. To isolate APOE genotype effects in a controlled human system, we use human induced pluripotent stem cell (hiPSC)-derived microglia from APOE4/4 parental clones that were CRIPSR/Cas9-corrected to APOE3/3, generating genetically matched APOE3/3-APOE4/4 isogenic clone pairs, differing only at the APOE locus (**Figure 7A; S7A–C)**, and differentiated into microglia over 42 days. Visual inspection of microglial morphology revealed APOE-dependent changes in cellular morphology, including alterations in cell body size and process complexity (**Figure 7B)**. Functional assessment using the pHrodo assay demonstrated that both APOE3 and APOE4 microglia are proficient in phagocytosis, with each genotype demonstrating the ability to effectively initiate and carry out phagocytic activity. (**Figure 7C; S7E)**.

**Figure 7:**
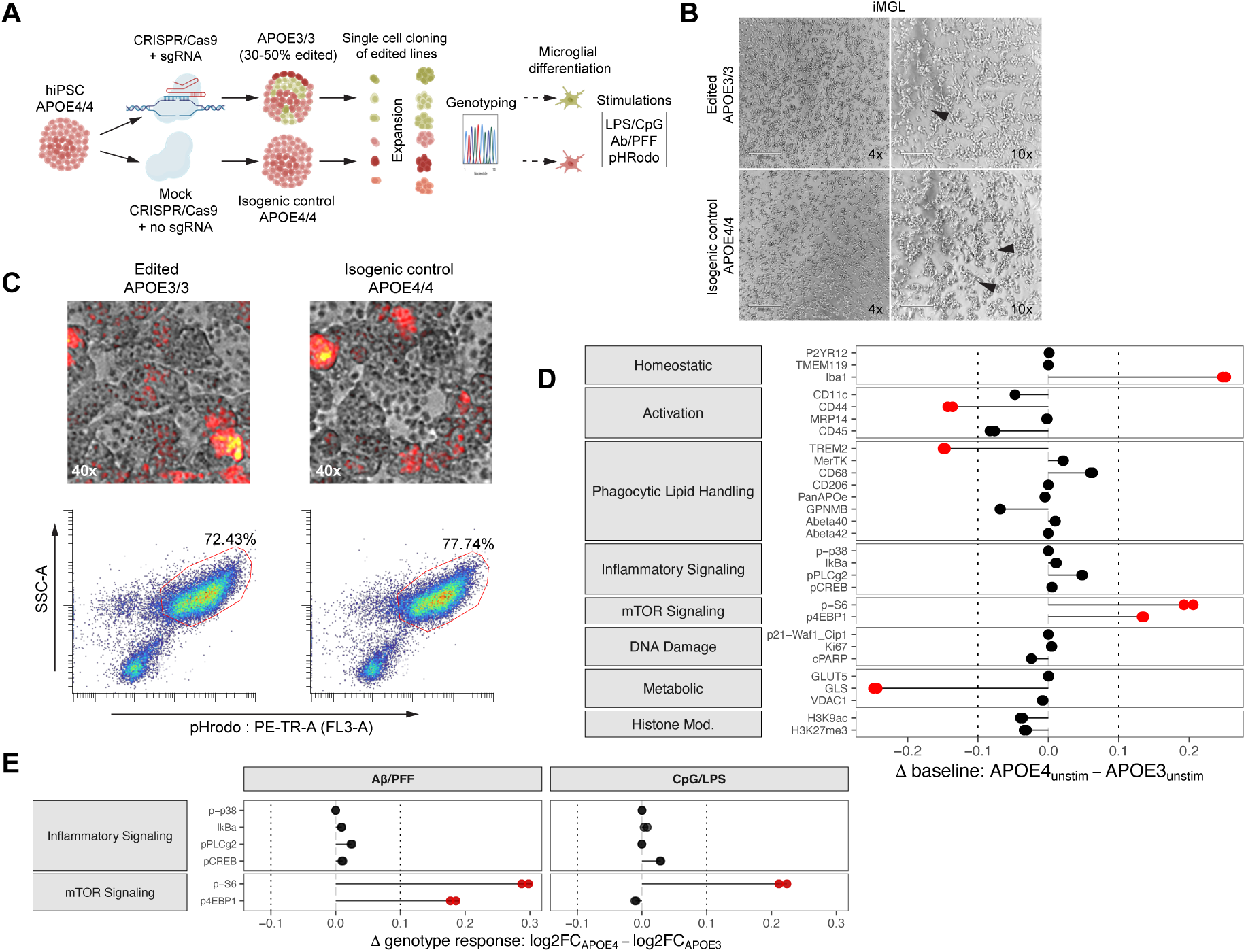
APOE shapes basal and stimulus-evoked cell states in isogenic iPSC-derived microglia. **(A)** Experimental design schematic. Human induced pluripotent stem cells (hiPSCs) carrying the APOE4/4 genotype were edited using CRISPR/Cas9 with sgRNA to generate APOE3/3 cells (30–50% editing efficiency). Mock-edited cells (CRISPR/Cas9 without sgRNA) were maintained as isogenic APOE4/4 controls. Single-cell cloning was performed to isolate individual edited and control clones, followed by clonal expansion and genotyping to confirm APOE status. Validated lines were differentiated into isogenic iPSC-derived microglia-like cells (iMGLs) differing exclusively at the APOE locus and subjected to functional stimulation assays (CpG/LPS, Aβ fibrils and tau pre-formed fibrils (Aβ/PFF), and pHrodo-based phagocytosis). iMGLs were profiled by CyTOF under three conditions: unstimulated, CpG/LPS stimulation, and Aβ/PFF stimulation. Two independent donor-derived line differentiations per genotype were analyzed. **(B)** Representative brightfield images of iMGLs derived from edited APOE3/3 and isogenic control APOE4/4 lines at 4× and 10× magnification. Both genotypes display comparable culture density following differentiation. **(C)** Phagocytic function assessed by pHrodo zymosan uptake after 1 hour. Representative images (40×) show internalized pHrodo-labeled zymosan particles (red) in edited APOE3/3 and isogenic control APOE4/4 iMGLs. Flow cytometry quantification (SSC-A vs pHrodo PE-TR-A) demonstrates the proportion of phagocytic cells, with comparable uptake between genotypes (percentages indicated). **(D)** Baseline differences in marker expression between APOE3 and APOE4 iPSC-derived microglia under unstimulated conditions. Expression values were pseudobulked by differentiation pairs and APOE genotype. For each differentiation pair, the genotype effect per marker was computed as the raw difference in pseudobulk expression (baseline_Δ_ = APOE4_unstim_ - APOE3_unstim_). Positive baseline_Δ_ values indicate higher expression in APOE4 microglia at baseline and vice versa. Points with genotype-associated shifts (|Δ| > 0.1) are highlighted in red. **(E)** APOE-dependent differences in stimulus response magnitude and directionality. Values represent the within-pair difference in response between genotypes (response_Δ_ = log_2_FC_APOE4_ - log_2_FC_APOE3_) for each marker and stimulus (CpG/LPS, Aβ/PFF). Positive values indicate a greater response in APOE4 microglia, whereas negative values indicate a greater response in APOE3 microglia. Points with genotype-associated shifts (|Δ| > 0.1) are highlighted in red.

At day 50, cells were treated with control vehicle (unstimulated), or acutely challenged with either CpG/LPS or Aβ fibrils and tau pre-formed fibrils (Aβ/PFF) (12 min stimulation), enabling assessment of both baseline genotype effects and stimulus-dependent responses. Cells were profiled by single-cell mass cytometry, as previously described in Bendall et al., 2012^108^ (**Figure 7D)**, with a panel spanning activation, phagocytic lipid handling, metabolism, and regulatory signaling activity **(Table S4)**. To quantify genotype-driven effects under unstimulated conditions, expression values were pseudobulked by matched differentiation pairs and APOE genotype, and a per-marker baseline difference was computed (baseline_Δ_ = APOE4_unstim_ – APOE3_unstim_). Notably, APOE4 microglia exhibited reduced CD44, TREM2, and GLS expression together with elevated mTOR signaling (pS6 and p4EBP1) and Iba1 expression, indicative of activation and structural remodeling (**Figure 7D; S7F)**. We next assessed early stimulus-induced responses following acute perturbation with either Aβ/PFF or CpG/LPS (**Figure 7E; S7G)**. Following CpG/LPS stimulation, APOE4 microglia mounted an amplified mTOR-linked response relative to isogenic APOE3/3 microglia, marked by increased pS6. Exposure to Aβ/PFF similarly induced elevated pS6 and p4EBP1 in APOE4/4 microglia, indicating enhanced activation of mTOR signaling in response to amyloid-related challenge.

Collectively, these findings suggest that APOE4 microglia are signaling-primed within a rejuvenated cellular context, exhibiting elevated basal and stimulus-induced mTOR pathway activity alongside reduced TREM2 and GLS expression – thus appearing to heighten acute activation potential while constraining lipid sensing and glutamine catabolic capacity relative to their isogenic APOE3 counterparts.

## DISCUSSION

Microglia occupy a continuum spanning homeostatic surveillance, transitional activation states, and dysfunctional phenotypes characterized by metabolic stress and loss of cellular identity. Relative to E3-AD, the microglial landscape in E4-AD is marked by a pronounced depletion of homeostatic populations and a concomitant expansion of APOE4-associated dysfunctional states that are distinct in both their cell-intrinsic molecular profiles and spatial niches. Among these states, we identify a previously under-appreciated microglial population that we termed Activation-Limited Microglia (ALMs). These cells are enriched in E4-AD brains and represent a non-canonical reactive state distinct from classical DAMs. Unlike DAMs, which accumulate near amyloid plaques and display robust phagocytic activation^11,12^, ALMs preferentially localize to regions characterized by metabolic insufficiency, astrocytic gliosis, and cellular senescence. At the molecular level, ALMs show reduced expression of lipid-handling markers ApoE and LAL, suggesting that they represent a hypometabolic, stress-adapted microglial state rather than a fully activated damage-response phenotype.

Importantly, these dysfunctional populations do not appear to arise *de novo*. Instead, our findings indicate that <u>APOE4 reshapes the trajectory through which microglia transition between functional</u> <u>states</u>, shifting the equilibrium away from homeostatic surveillance and toward stress-associated outcomes. Trajectory inference indicates that genotype-dependent differences in microglial behavior are modulated through selective divergence during intermediate stages of state progression. The earliest lineage bifurcation is notable not simply for the persistence of core microglial identity and inflammatory features in Reactive 2 at late pseudotime, but also the elevated expression of HLA-DR, GPNMB, and CD68 at the branch point itself. This suggests that lineage divergence may be initiated during the immediate response to acute inflammatory cues^61,109–112^, rather than reflecting differential resolution of inflammation at later stages.

One parsimonious interpretation is that microglial trajectories differ not primarily in inflammatory exposure, but in how that stimulus is processed over time. Cells along the early-divergence lineage may enter a primed inflammatory state, conferring a broadly accessible reactive capacity under chronic neurodegenerative stress. In contrast, the later-diverging lineages, terminating in E4-AD-enriched ALM and dystrophic microglial states, may reflect alternative outcomes that arise when microglia fail to stabilize homeostatic and canonical damage-response programs. Attenuation of TREM2 expression in these latter lineages post-branching aligns with prior work implicating the TREM2-APOE axis in supporting full DAM conversion under chronic demyelination stress, and that disruption of this axis inhibits maintenance of microglial metabolic fitness^10^. Rather than invoking a singular APOE4-specific defect, our data supports a model in which APOE biases the routing of microglia through these trajectories following acute inflammation, shaping whether cells retain immune competence or transition toward hypo-responsive or dystrophic phenotypes.

This trajectory-based framework helps reconcile seemingly conflicting descriptions of APOE4 microglia reported in the literature. APOE4 microglia have been alternately described as persistently hyper-inflammatory^13–15^ or as dysfunctional and hypo-responsive^20,21,24^, interpretations that are often framed as competing models. <u>Rather than representing mutually exclusive states, our findings suggest that</u> <u>these observations may instead reflect different phases along an APOE4-biased microglial fate</u> <u>progression that progressively erodes homeostatic surveillance during neurodegeneration.</u> Consistent with this proposal, we found that iPSC-derived APOE4 microglia display heightened basal activation, including increased mTOR pathway activity. While early-life hyper-reactivity may initially enhance immune responsiveness, it also predisposes APOE4 microglia to chronic inflammatory signaling. The late-stage APOE4 microglial phenotypes observed in disease-aged human tissue may reflect the consequence of persistent, unresolved activation that culminates in functionally constrained, exhaustion-like states. In this respect, APOE4 microglia may be conceptually analogous to exhausted T cells in chronic infection or cancer, where prolonged engagement of inflammatory signaling programs ultimately erodes their functional capacity^113,114^. This phenomenon mirrors the maladaptive state of APOE4 microglia described in this work, in which features of activation are retained, yet execution of core effector programs is impaired^115^. Interestingly, APOE4-associated microglia exhibit elevated Ikaros (IKZF1) motif accessibility, an epigenetic regulator previously identified as a key driver of T cell exhaustion via silencing of effector genes under persistent stimulation^116^. Consistent with this, APOE4 microglia also upregulated *NFKBIZ*, a regulator of effector-like transitory exhausted T cell differentiation^117^, further suggesting convergence on a conserved chronic stress-adaptation program.

This work has important implications for understanding APOE4-driven neurodegeneration. Therapeutic strategies aimed solely at suppressing inflammation in late-stage disease may overlook the underlying metabolic vulnerabilities of APOE4 microglia. Instead, strategies that stabilize microglial metabolic programs, support lysosomal function, and preserve homeostatic identity may prove critical for maintaining microglial competence in APOE4 homozygotes.

## LIMITATIONS OF THE STUDY

These findings are derived from a rigorously curated human cohort with genotype-defined groups, low post-mortem interval, strict neuropathological inclusion criteria, and exclusion of major neurodegenerative co-pathologies, enabling high-fidelity characterization of APOE-dependent microglial states in a controlled disease context. While both male and female donors were included, the study was designed to resolve genotype- and disease-associated effects and is not powered to systematically assess sex-specific differences. As emerging evidence suggests that APOE4-associated risk and microglial responses may be sex-dependent, future studies will be required to define how these variables intersect with the trajectory-based framework described here. An important consideration in this study is that trajectory inference was performed on cross-sectional imaging data from human autopsy tissue. As such, the inferred lineages are better understood as a data-driven approximation of underlying biological continua, rather than direct measurements of temporal progression or definitive developmental paths. Definitively establishing the temporal ordering and causal relationships amongst these microglial states will require complementary experimental systems such as perturbation studies and longitudinal sampling in model systems that allow for controlled manipulation of APOE genotype. However, as the phenotypes observed in end-stage disease may emerge only over prolonged timescales, fully recapitulating the relevant temporal dynamics experimentally may not be feasible.

## ACKNOWLEDGEMENTS

This work was supported by National Institutes of Health grants U01AG072573, P01AG073082, P30AG066515, R01AG056287, R01AG057915, R01AG068279, U19 AG065156, U24CA224309, P30AG066515, U54HL165445, R01AG078702, R01AG088656, R01NS121404, P01AG036695, R01CA251858, R01CA240638, and T32AI007290. R.E. is supported by National Science Scholarship, Agency for Science, Technology, and Research (A*STAR), Singapore.

We are grateful to the Banner Sun Health Research Institute Brain and Body Donation Program of Sun City, Arizona for the provision of human biological materials. The Brain and Body Donation Program has been supported by the National Institute of Neurological Disorders and Stroke (U24 NS072026 National Brain and Tissue Resource for Parkinson’s Disease and Related Disorders), the National Institute on Aging (P30 AG019610 and P30 AG072980, Arizona Alzheimer’s Disease Center), the Arizona Department of Health Services (contract 211002, Arizona Alzheimer’s Research Center), the Arizona Biomedical Research Commission (contracts 4001, 0011, 05-901 and 1001 to the Arizona Parkinson’s Disease Consortium) and the Michael J. Fox Foundation for Parkinson’s Research. PBMC samples for iPSC derivation were obtained from the Stanford Alzheimer’s Disease Research Center, NIH/NIA grant P30 AG066515.

## METHODS

### I. Human brain tissue

Human formalin-fixed paraffin-embedded (FFPE) brain samples were selected from the Brain and Body Donation Program at Banner Sun Health Research Institute, (brainandbodydonationprogram.org)^118^. Inclusion criteria were homozygosity for one of the *APOE* alleles, annual clinical assessment of probable AD dementia with high level for AD neuropathologic change or annual clinical assessment of normal cognition with no or low AD neuropathologic change^32,33^. Exclusion criteria were clinical diagnosis of neurological disease other than AD and any pathologic evidence of neurodegenerative co-morbidity from Lewy body disease, glial cytoplasmic inclusion, pathologic TDP-43 inclusion, hippocampal sclerosis, or any other less common neurodegenerative disease. All cases and controls satisfying these criteria with available tissue blocks were analyzed. The resulting study cohort (n = 12) comprised 4 groups: *APOE* ε3/3 individuals with AD dementia (E3-AD), *APOE* ε4/ε4 individuals with AD dementia (E4-AD), *APOE* ε3/3 cognitively normal individuals (E3-CN), and *APOE4* ε4/ε4 cognitively normal individuals (E4-CN). Donor details are listed in Table S1.

### II. Method Details

#### Antibody preparation

Antibodies were first screened by immunohistochemistry (IHC) on FFPE brain sections to select for the best performing clones. These were then conjugated to isotopic metal reporters as described previously. Following conjugation antibodies were diluted in Candor PBS Antibody Stabilization solution (Candor Bioscience). Antibodies were either stored at 4°C or lyophilized in 100 mM d-(+)-trehalose dehydrate (Sigma-Aldrich) with ultrapure distilled H2O for storage at −20 °C. Before staining, lyophilized antibodies were reconstituted in a buffer of Tris (Thermo Fisher Scientific), sodium azide (Sigma-Aldrich), ultrapure water (Thermo Fisher Scientific) and antibody stabilizer (Candor Bioscience) to a concentration of 0.05 mg ml−1. The antibodies, metal reporters and staining concentrations are listed in Extended Data Table S2. For detailed metal-antibody protocol MIBItag see Bosse et al., 2021^119^.

#### Tissue staining

FFPE brain tissues were sectioned (5-μm section thickness) from tissue blocks on gold and tantalum-sputtered microscope slides. Slides were baked at 70 °C overnight, followed by deparaffinization and rehydration with washes in xylene (3×), 100% ethanol (2×), 95% ethanol (2×), 80% ethanol (1×), 70% ethanol (1×) and ddH2O with a Leica ST4020 Linear Stainer (Leica Biosystems). Slides next underwent antigen retrieval by submerging sides in 3-in-1 Target Retrieval Solution (pH 9, DAKO Agilent) and incubating at 97 °C for 40 min in a Lab Vision PT Module (Thermo Fisher Scientific). After cooling to room temperature for 1 h, slides were washed in wash buffer (1× PBS IHC Washer Buffer with Tween 20 (Cell Marque) with 0.1% (w/v) bovine serum albumin (Thermo Fisher)). Next, all slides underwent two rounds of blocking, the first to block endogenous biotin and avidin with an Avidin/Biotin Blocking kit (BioLegend). Slides were then washed with wash buffer and blocked for 1 h at room temperature with 1× TBS IHC Wash Buffer with Tween 20 with 3% (v/v) normal donkey serum (Sigma-Aldrich), 0.1% (v/v) cold fish skin gelatin (Sigma-Aldrich), 0.1% (v/v) Triton X-100 and 0.05% (v/v) sodium azide. The first round of staining was carried out with free indium 1153+ (8 mM diluted in PBS, Fluidigm) in staining buffer (1× TBS IHC Wash Buffer with Tween 20 with 3% (v/v) normal donkey serum) and incubated overnight at 4°C in a humidity chamber. The following day, slides were washed twice for 5 min on a shaker in wash buffer. The second round of staining was conducted using the cocktail of metal-conjugated antibodies prepared in a staining buffer at their respective concentrations and filtered through a 0.1-μm centrifugal filter (Millipore) before incubation with tissue overnight at 4°C in a humidity chamber. Following the overnight incubation with the antibody cocktail, slides were washed twice for 5 min in wash buffer. On the third day, anti-biotin 152 Eu was prepared as described and incubated with the tissues for 1 hour at 4°C in a humidity chamber. Following staining, slides were washed twice for 5 min in wash buffer and fixed in a solution of 2% glutaraldehyde (Electron Microscopy Sciences) solution in low-barium PBS for 5 min. Slides were then washed in PBS (1×), 0.1 M Tris at pH 8.5 (3×) and ddH2O (2×) and then dehydrated by washing in 70% ethanol (1×), 80% ethanol (1×), 95% ethanol (2×) and 100% ethanol (2×). Slides were dried under vacuum overnight before imaging. For detailed slide staining protocol for MIBI-TOF imaging see Bosse et al., 2021^120^.

#### Immunohistochemistry (IHC)

For IHC screening of panel antibodies, FFPE human hippocampal tissue was sectioned onto standard glass slides at 5-μm thickness. Slides containing tissue were baked at 70 °C overnight. Tissue sections were then processed and stained in a Sequenza staining rack (Thermo Fisher) with a single primary antibody. The IHC protocol mirrors the MIBI protocol, with the addition of blocking endogenous peroxidase activity with 3% (v/v) H2O2 (Sigma-Aldrich) in ddH2O after epitope retrieval. On the second day of staining, instead of proceeding with the MIBI protocol, tissues were washed twice for 5 min in wash buffer and stained using ImmPRESS universal (anti-mouse/anti-rabbit horseradish peroxidase) kit (Vector Laboratories). For detailed IHC staining techniques see Bosse et al., 2021^121^.

#### Image acquisition on MIBI-TOF

Multiplexed ion beam imaging by time-of-flight (MIBI-TOF) was performed on a commercial MIBIscope instrument (M13) equipped with a Hyperion ion source (Ionpath). Slides were sputter-coated with a 10 nm layer of gold prior to imaging to improve sample conductivity and facilitate dissipation of accumulated surface charge. Image acquisition was performed using a sequential pre-raster and acquisition strategy. Pre-rastering was conducted in fine mode (40nA beam current; 400μm × 400μm FOV; 512 × 512 pixels; 0.25ms dwell time per pixel) to remove the gold layer from regions of interest. During pre-rastering, a 2000 V sample bias was applied, and secondary ions were not funneled into the TOF chamber to preserve detector integrity. Following gold removal, final image acquisition was performed in coarse mode using a 400μm × 400μm FOV at 1024 × 1024-pixel resolution with a 0.5ms dwell time per pixel (40nA beam current). During acquisition, secondary ions were funneled into the TOF chamber for detection, and detector voltage was set to the autoGain value determined from the preceding acquisition run. To reduce cumulative charging artifacts during tiled imaging, FOV acquisition order was randomized. PMMA standards were used for routine quality control under coarse acquisition settings (200μm × 200μm; 512 × 512 pixels; 0.5ms dwell time).

#### Low-level image processing

Background signal contamination was removed via the Rosetta algorithm, which uses a flow-cytometry style compensation approach to remove spurious signals. Images were then normalized to ensure consistent intensity across each run. All image processing tools in the MIBIScope-processing toffy toolkit can be accessed at the following repository: https://github.com/angelolab/toffy.

#### Single-object segmentation

Single-object segmentation of multiplexed TIFF images was performed using ezSegmenter, an intensity-thresholding based tool. Object composite channels were created using combined lineage phenotyping marker expression **(Table S5)**. Composite channels were then Gaussian blurred and thresholded on either a given fixed value, or an adaptive thresholding method, to generate object masks. Parameters tuned include standard deviation for Gaussian kernel, minimum and maximum object size (in pixels), and per-FOV-percentile threshold value (integer) for image thresholding. Masks are used to extract pixel-level signal intensities across each channel on a per-object basis, which are then size normalized before output into a cell table csv file. Single cell or object data was then imported into R for cube root transformation and 99.9 percentile normalization. ezSegmenter is available at https://github.com/angelolab/ark-analysis.

#### Extracting spatial features from multiplexed TIFFs

Spatial features were extracted from multiplexed TIFF images to characterize the local tissue microenvironment surrounding individual microglial cells. All features were computed relative to segmented microglial cells and their immediate microenvironment. For each microglial cell, a 20-pixel expansion zone was generated by morphologically dilating the original cell segmentation mask, thereby preserving the shape of the primary cell while extending its boundary uniformly outward. This approach maintains native cell geometry and avoids directional bias that can arise from circular or centroid-based neighborhood definitions.

##### Pixel-based spatial features

Mean marker intensity was calculated within expansion zones on a per-marker basis, yielding pixel-level measures of local protein abundance in the immediate pericellular environment. Binary spatial features corresponding to myelinated regions, astrocyte gliosis, and free iron-rich zones were derived using marker-specific intensity thresholds. For each feature, thresholds were selected based on the distribution of marker signal within microglial neighborhoods: Fe signal for iron-associated niches, CD44 and GFAP co-expression for gliosis-associated niches, and CNPase signal for myelin-rich niches. Thresholds were validated through manual annotation of a sub-sample of fields-of-view (FOVs) classified as positive or negative for each feature, confirming that selected threshold values indeed maximized separation between annotated classes. These thresholds were subsequently applied across all images to assign feature-association positivity within each microglial expansion zone.

##### Object-based spatial features

Object-based spatial features were computed by identifying secondary segmented objects within each microglial expansion zone, including PHF-tau^+^ objects, γH2A.X^+^ objects, and amyloid-β plaques. For each interaction between a microglial cell and a secondary object within the expansion zone, the following attributes were recorded: (i) the unique identifier of the secondary object, (ii) the x and y centroid coordinates of the secondary object, (iii) the percentage of the secondary object’s area overlapping with the expansion zone, and (iv) the minimum Euclidean distance (in pixels) between the secondary object and the segmentation boundary of the primary microglial cell. Together, these measurements captured both the spatial proximity and degree of interaction between microglia and pathological or stress-associated features in the surrounding tissue microenvironment.

#### Calculating microglial plaque coverage and minimum distance to plaques

To quantify microglial coverage around plaques, we calculated the expected area of microglia in the expansion region assuming uniform distribution across the field of view (FOV). We then compared this to the observed microglial area within each plaque’s 50-pixel (19.5µm) expansion zone. The resulting enrichment ratio – defined as the log_2_ ratio of observed to expected microglial area coverage – served as a measure of microglial recruitment to plaques. Positive values indicate local enrichment, while negative values reflect under-representation relative to a uniform spatial distribution. This metric allows for direct comparison across clusters or plaque types while accounting for baseline microglial density in the tissue. For each microglial cell within a given plaque’s expansion region, we defined its distance to plaque pathology as the minimum number of pixels between the cell’s and plaque object’s segmentation mask.

#### RNA preprocessing

Low-quality cells were excluded based on standard quality control metrics, retaining cells with >1,000 detected transcripts, >200 genes, and <5% mitochondrial and ribosomal read content. Contaminating macrophages and cycling cells were additionally removed based on canonical marker expression. Gene expression counts were log-normalized using *Seurat*^122^ (v5.2.1), and the resulting normalized expression matrix was extracted for downstream analyses, including use as the response matrix in linear modeling.

#### ATAC preprocessing and peak calling

Arrow files were loaded into *ArchR*^123^ (v1.0.3) for downstream analysis. Quality control filtering retained cells with transcription start site (TSS) enrichment ≥ 4 and ≥ 1,000 unique fragments, and inferred doublets were removed. Microglia were identified based on paired RNA transcript annotations. Dimensionality reduction was performed using iterative latent semantic indexing (LSI) via addIterativeLSI with four iterations, and cells were clustered using addClusters with default parameters. Pseudobulk replicates were generated with addGroupCoverages, and a consensus peak set was called using MACS3 via addReproduciblePeakSet with default parameters, yielding a unified peak-by-cell matrix for downstream analyses.

#### PBMCs and iPSC reprogramming

PBMCs were received directly from the Stanford Alzheimer’s Disease Research Center (ADRC) PBMCs were collected in a Ficoll density gradient separation tube. Reprogramming was performed by AlStem Services. The parental cell lines were reprogrammed into iPSCs from PBMCs via transfection with non-integrating episomal or mRNA vectors encoding OCT4, SOX2, KLF4, and L-MYC. Colonies were selected based on morphology. Clone quality was confirmed by mycoplasma, and sterility reports confirmed that clones were free of bacterial and fungal contamination.

Pluripotency for all clones was confirmed using characterization assays, including immunostaining for TRA-1-60 and OCT4, and the assessment of alkaline phosphatase (AP) activity. These assays were performed following initial clone picking to confirm pluripotency.

#### Human induced pluripotent stem cell (hiPSC) derivation, culture, and maintenance

The hiPSC lines (APOE 3/3, n = 4; APOE 4/4, n = 4), derived from two donors (four clones per donor), were maintained under feeder-free conditions using growth factor-reduced Matrigel (1:30 dilution, Corning) in complete STEMflex medium (cSFM; Thermo Scientific) in a humidified incubator (5% CO2, 37°C). Cells were fed daily with fresh medium and passaged every 3–5 days at a 1:8–1:10 ratio using ReLeSR (STEMCELL Technologies) according to the manufacturer’s instructions.

#### CRISPR derivation and validation of APOE R130C iPSC knock-in clones

Human iPSC lines were used to generate APOE R130C knock-in clones carrying the c. CGC>TGC nucleotide substitution, corresponding to the R130C amino-acid change in APOE. The engineered clones were produced as iPSC knock-in clones by EditCo Bio. The targeted gene was APOE, using transcript ENST00000252486, with the guide RNA cut site reported at chr19:44,908,683. The guide RNA sequence used for targeting was GCGGACAUGGAGGACGUGCG. Homology-directed repair was supported by a donor repair sequence containing the intended R130C edit. Following CRISPR-mediated editing and clonal expansion of three edited iPSC clones were selected for delivery and quality-control analysis. All three clones were reported as homozygous APOE R130C knock-in clones and were cryopreserved. Mycoplasma testing was negative for all clones. Genotype confirmation was performed by targeted next-generation sequencing of the APOE locus using primers flanking the edited region. The primer sequences were forward primer 5′-CTGGAGGAACAACTGACCCC-3′ and reverse primer 5′-GGCGCTTCTGCAGGTCAT-3′.

#### Differentiation of microglia-like cells (iMGLs) from hiPSCs

To generate induced microglia-like cells (iMGLs), hiPSCs were first differentiated into hematopoietic progenitor cells (iHPCs) for 12 days, followed by 28 days of microglial differentiation using previously established protocols by McQuade et al., 2018^124^ and McQuade and Blurton-Jones, 2021^125^. Briefly, hiPSCs were cultured to ∼70% confluence and passaged as small clumps (100–200 µm) onto fresh Matrigel (1:90 dilution) in cSFM at low density (1:150–1:200) using ReLeSR. Differentiation into iHPCs began the following day using the StemDiff Hematopoietic Kit (STEMCELL Technologies) for 10–12 days, according to the manufacturer’s instructions. On days 10–12, cells were harvested and validated by flow cytometry for CD34⁺, CD45⁺, and CD43⁺ iHPC populations. Microglial specification was induced using a cytokine cocktail consisting of IL-34 (100 ng/mL; PeproTech), TGF-β1 (50 ng/mL; PeproTech), and M-CSF (25 ng/mL; PeproTech) for 25 days. iMGL maturation was subsequently achieved by supplementing the medium with CD200/OX2 (100 ng/mL; Abclonal) and CX3CL1 (100 ng/mL; BioLegend) for an additional 3 days. To induce a pro-inflammatory state, iMGLs were cultured for an additional 2–3 days in medium lacking the homeostatic factors TGF-β1, CD200, and CX3CL1. The basal iMGL differentiation medium consisted of phenol red–free DMEM/F12 (1:1), insulin–transferrin–selenite (2X; Thermo Scientific), N2 supplement (0.5X; Thermo Scientific), B27 supplement (2X; Thermo Scientific), monothioglycerol (400 µM; Sigma-Aldrich), GlutaMAX (1X; Thermo Scientific), non-essential amino acids (1X; Sigma-Aldrich), and insulin (5 µg/mL; Sigma-Aldrich).

#### iMGL stimulation and phRodo phagocytosis assay

For stimulation assays, pro-inflammatory state iMGLs were challenged with lipopolysaccharide (LPS, Invivogen), CpG oligodeoxynucleotides (Fisher Scientific), human amyloid-β pyroglutamate 3–42 pre-formed fibrils (StressMarq), and active human recombinant tau (K18) P301L mutant pre-formed fibrils (StressMarq) using previously established protocols^126,127^. Briefly, 1.0 × 10⁶ iMGLs per condition were collected, counted, and resuspended in stimulation medium (basal iMGL differentiation medium supplemented with 2 mM EDTA, heparin, and benzonase) to a final volume of 800 µL per tube. Cells were distributed into three conditions: unstimulated control, inflammatory stimulation (LPS + CpG), and neurodegenerative stimulation (amyloid-β fibrils + tau pre-formed fibrils). A 100× stimulation solution (200 µL) was added to each tube to reach a final volume of 1mL. Final concentrations were 100 ng/mL LPS and 2 µg/mL CpG for inflammatory stimuli, and 1µM amyloid-β fibrils and 1µM tau pre-formed fibrils for neurodegenerative stimuli. Cells were stimulated at 37 °C and collected at defined time points (0 and 12 min). Reactions were terminated by direct addition of 16% paraformaldehyde (100µL per 1mL), followed by incubation for 10 min at room temperature. Cells were then washed with cell staining medium, pelleted by centrifugation, and stored at −80 °C until downstream CyTOF barcoding and analysis.

For phagocytosis assays, pro-inflammatory state iMGLs were seeded at a density of 5 × 10⁵ cells onto vitronectin-coated plates and allowed to adhere for 2 days. Cells were then incubated with pHrodo™ Red Zymosan Bioparticles™ Conjugate (0.5 mg/mL; Thermo Fisher Scientific) for 1 h at 37 °C. Live-cell imaging was performed every 20 min using the ImageXpress Pico (Molecular Devices). Following incubation, cells were collected and fixed for CyTOF analysis as described above. Single-cell suspensions for CyTOF were barcoded and stained as previously described by Amouzgar et al., 2025^128^. Cells were stained with a panel of phenotypic, metabolic, and phosphorylation-specific antibodies. Cells were acquired on a CyTOF2 mass cytometer (Fluidigm), and data were analyzed using CellEngine (CellCarta).

### III. Statistical Analysis

#### Global co-expression network construction

To characterize global inter-marker relationships across the multiplexed imaging dataset, we constructed a co-expression network using all segmented objects, including microglia, neurons, astrocytes, tau- and amyloid-positive structures, and vascular elements. Marker intensities were log-transformed and Z-scored per marker prior to analysis. Pairwise Spearman correlation coefficients were computed across all markers using the R package *Hmisc* (v5.1-1), yielding correlation coefficients (ρ) and associated p-values for each marker pair. Multiple testing correction was performed using the Benjamini-Hochberg procedure. Marker pairs were retained as network edges if they met both effect size and significance thresholds (|ρ| ≥ 0.3 and FDR < 0.05), and the resulting filtered correlation matrix was used to define the network adjacency structure. Network construction and analysis were performed using the R package *igraph* (v2.0.1.1). Node strength, defined as the sum of absolute correlation weights, was computed, and used for node scaling.

#### Linear mixed-effects modeling and planned marginal means comparisons

Single-cell continuous features were analyzed across multiple brain cell types (astrocytes, microglia, and neurons) using linear mixed-effects models. For each feature, outcomes were standardized (z-scored) prior to modeling. Categorical predictors were coded with APOE3/3 as the reference genotype and cognitively normal (CN) as the reference diagnosis. For each marker, a single cell-level mixed-effects model was fit:

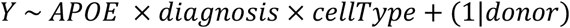

where Y denotes the standardized feature value. Models were estimated using restricted maximum likelihood (REML) with the *lme4* framework, and fixed-effect inference was obtained using lmerTest. A random intercept for donor (caseID) was included to account for within-donor correlation and mitigate pseudoreplication. Planned contrasts were evaluated using estimated marginal means (*emmeans*), with equal weighting across conditioning factor levels (weights = “equal”) to ensure design-balanced comparisons. All contrasts were performed separately within each cell type. Genotype-driven effects were assessed by comparing E4 versus E3, averaged over diagnosis, using marginal means of APOE genotype within cell type. Diagnosis-driven effects were assessed by comparing AD versus CN, averaged over genotype, within each cell type. Genotype-by-diagnosis interactions were tested using a difference-in-differences contrast: (E4, AD−E4, CN)−(E3, AD−E3, CN), computed from marginal means of the full APOE genotype × diagnosis interaction. For diagnosis-stratified analyses, diagnosis effects (AD − CN) were additionally estimated within each genotype (E3 and E4) and cell type.

#### Computation of condition-associated relative likelihood estimates

To quantify the combinatorial effect of both APOE genotype and diagnostic status on hippocampal microglial single-cell states, we utilized a graph signal processing approach, MELD (Manifold Enhancement of Latent Dimensions^47^, that infers the relative likelihood estimate of observing a given cell in one of four APOE-diagnosis conditions, thereby allowing us to map a continuous “APOE-diagnosis effect” across the proteomic manifold space. Briefly, MELD learns a low-dimensional manifold of the proteomic space and constructs a cell similarity graph where each cell is represented by a single node in the graph. Computing the kernel density estimate (KDE) over the cell similarity graph using graph signals representing indicator vectors for each APOE-diagnosis condition allows us to calculate the condition-associated density estimate (i.e., the density of cells associated with each condition over the manifold). The relative condition-associated likelihood is calculated via Bayes-style normalization, yielding a posterior probability for each cell conditioned on its manifold neighborhood. For the paired snRNA-seq and snATAC-seq data, MELD was applied on the k-nearest neighbors (kNN) graph generated from the learned MultiVI latent components. Analysis was performed in python using the *meld* package (v1.0.2).

#### Notation for MELD-derived condition-associated likelihood estimates

Throughout this study, we refer to MELD-derived relative likelihood estimates of a cell’s association with a specific APOE-diagnosis condition using the notation *p*(condition). For example, *p*(E4-AD) denotes the relative likelihood that a given cell is associated with the APOE4 Alzheimer’s disease group, as inferred by MELD. Similarly, *p*(E3-CN), *p*(E3-AD), and *p*(E4-CN) refer to likelihood scores for the APOE3 cognitively normal control, APOE3 Alzheimer’s disease, and APOE4 cognitively normal control groups, respectively. These estimates reflect the extent to which a cell’s molecular profile aligns with a given condition and are used throughout the manuscript to identify and interpret disease-associated cellular states.

#### Gaussian Mixture Model (GMM)-based clustering of hippocampal microglia

Unsupervised clustering of hippocampal microglia was performed using a GMM framework implemented in the *mclust* R package (v6.0.1). GMMs model the data as a mixture of multivariate Gaussian distributions, enabling probabilistic assignment of cells to clusters while accommodating overlapping and transitional phenotypes. The optimal number of mixture components was determined by fitting models with varying numbers of components and evaluating model fit using both the Akaike Information Criterion (AIC) and the Bayesian Information Criterion (BIC). An eight-component solution was selected based on concordant support from both criteria and biological interpretability. For downstream analyses, cells were hard assigned to clusters based on their maximum posterior membership probability. Cluster identities were subsequently annotated using canonical microglial markers and morphological features.

#### Differential protein expression analysis between clusters

Differential expression analyses between pairs of microglial clusters were performed using a non-parametric framework. For each marker, differences in expression between clusters were assessed using the Wilcoxon rank-sum test, yielding two-sided p-values. To account for multiple testing across markers, p-values were adjusted using the Benjamini-Hochberg (BH) procedure. Effect sizes were quantified using Cliff’s delta, calculated with the *effsize* R package (v0.8.1), providing a robust, distribution-free measure of the magnitude and direction of expression differences. Markers were considered differentially expressed if they met both statistical and effect-size criteria: BH adjusted p < 0.05 and |Cliff’s delta| ≥ 0.33, corresponding to at least a medium effect size.

#### Spatial feature association by cluster

To determine whether the frequency of feature-associated microglia differed across clusters, we annotated each microglial cell as ‘feature-associated’ or ‘non-feature-associated’ based on expansion-zone expression of relevant markers **(Table S6)**. Proportions of feature-associated cells were first summarized for each cluster. A chi-square test of independence was applied to evaluate whether the distribution of feature association differed across clusters. To estimate effect sizes and compare clusters pairwise, we fit a logistic regression model with feature association (yes/no) as the dependent variable and cluster identity as the predictor. Estimated probabilities and odds ratios with 95% confidence intervals were derived using the *emmeans* package in R (v1.10.2). Multiple comparisons were adjusted using the Benjamini-Hochberg (BH) procedure.

#### Quantifying local microenvironmental diversity

To quantify the compositional diversity of the local microenvironment surrounding individual microglial cells, we computed Shannon entropy over the distribution of neighboring object types. Neighbor data were represented in long format, with each row corresponding to a single neighboring object assigned to a primary microglial cell and annotated by its object class. For each microglial cell, the number of neighboring objects belonging to each class (*n_t_*) was counted, and the total number of neighbors was defined as *N* = ∑*_t_ n_t_*.

Class proportions were calculated as:

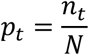

Shannon entropy was then computed using base-2 logarithms:

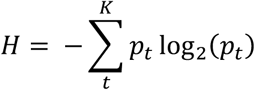

where higher values indicate greater heterogeneity in neighbor composition. To enable comparisons across cells with differing numbers of observed neighbor types, entropy values were normalized by the maximum possible entropy:

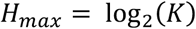

where K denotes the number of distinct neighbor types. By default, K was defined on a per-cell basis as the number of unique neighbor classes observed for that cell; alternatively, a fixed global K was used when the full set of possible neighbor types was known.

#### Pairwise co-occurrence analysis of spatial niche features

Pairwise co-occurrence of a predefined set of spatial features was quantified using binary feature annotations indicating presence or absence per cell (e.g., white matter association, astrocytic gliosis, tau-positive neighbors, amyloid plaques, senescent neighbors, and free iron-rich zones). Analyses were performed both globally across all cells and stratified by microglial cluster. For each pair of features (*f*_1_, *f*_2_), we constructed a 2×2 contingency table representing the joint distribution of feature positivity across all observations: *n*_11_(both positive), *n*_10_ (*f*_1_ positive only), *n*_01_ (*f*_2_ positive only), and *n*_00_ (both negative). Statistical significance of co-occurrence was assessed using a two-sided Fisher’s exact test, yielding an odds ratio (OR) and p-value for each feature pair. To ensure stable effect size estimates in sparse tables, ORs were computed using a Haldane-Anscombe continuity correction (addition of 0.5 to all table entries) and reported on a log_2_ scale (log_2_(OR)) to symmetrize enrichment (positive values) and mutual exclusivity (negative values). Multiple testing correction was performed using the Benjamini-Hochberg (BH) procedure.

#### Between-cluster comparison of feature co-occurrence

For each pair of binary spatial features (e.g., white matter vs. gliosis), we first computed per-group 2×2 counts within the cells of interest: n_11_ (both present), n_10_, n_01_, and n_00_. From these we derived a continuity-corrected odds ratio (Haldane–Anscombe, +0.5 in each cell) and its strength on a log_2_ scale (log_2_OR); Jaccard overlap was also calculated for visualization. To test whether the association between the two features differed between two groups (e.g., Cluster A vs. Cluster B), we constructed a 2×2×2 contingency table with factors Feature A (0/1), Feature B (0/1), and Group (A/B) and fit Poisson log-linear models with and without the three-way interaction term (A×B×Group). A likelihood-ratio test of this interaction provides a p-value for heterogeneity of the odds ratios (i.e., whether co-occurrence strength differs between groups; equivalent to a homogeneity-of-ORs test). We report Δlog_2_OR = log_2_ OR_B_ – log_2_OR_A_ alongside the p-value and control the false discovery rate across all tested feature pairs using Benjamini–Hochberg. Rare features (<5% prevalence within a group) were excluded to avoid unstable estimates; within-group enrichment edges were additionally filtered to retain positive associations (log_2_OR > 0) that were significant after FDR correction.

#### Random Forest classification

Cluster identity was modeled using a random forest classifier implemented with the *ranger* package (v0.16.0) in R. All pixel- and object-level spatial features were included as predictors, with the categorical cluster label as the outcome. To address the substantial class imbalance in the dataset (where majority of cells belong to the homeostatic clusters), we implemented inverse-frequency class weights such that misclassification of rare clusters was subject to a heavier penalty than that of abundant clusters. Model performance was assessed using stratified K-fold cross-validation, with folds constructed so that every cluster was represented in each fold. Evaluation metrics included overall out-of-bag error, per-class precision, recall and F1-scores, as well as macro-averaged metrics that weight all clusters equally. To further quantify predicted performance under imbalance, we computed class-specific receiver operating characteristic (ROC) and precision-recall (PR) curves with their corresponding area under the curve (AUC) values. To assess feature importance, we employed permutation importance, which estimates the decrease in predictive performance when the values of a feature are randomly permuted, thereby disrupting its relationship with the outcome.

#### Mixed-effects ordinal trend testing across ordered microglial states

To test for stepwise changes in marker expression across the inferred continuum (Homeostatic → Transitional → Early Dystrophic → Dystrophic), we performed a mixed-effects ordinal trend analysis. Analyses were restricted to nucleated microglia from the four ordered states. For chromatin-associated readouts, markers were normalized to total Histone H3 signal per cell. For each marker, microglial state was encoded as an ordered factor and state progression was modeled using an ordinal numeric variable. Linear mixed-effects models with random intercept for donor to account for within-donor correlation were fit using *lme4* (v1.1-35.1), and fixed-effect inference (including degrees of freedom) was obtained using *lmerTest* (v3.1-3) with Satterthwaite approximation. The primary trend statistic was the fixed-effect slope, reported as the standardized per-step effect size, with 95% confidence intervals extracted from the fitted model. Model fit was summarized using marginal and conditional *R*^2^ values.

#### Trajectory inference and pseudotime analysis

Trajectory inference was performed with the *slingshot* package in R^84^ (v2.10.0), which reconstructs lineage relationships and estimates continuous pseudotime from low-dimensional embeddings. Principal component analysis (PCA) was performed on a curated panel of microglial markers to generate the low-dimensional representation used for trajectory inference. The first four principal components were retained as input for trajectory inference. Categorical cluster labels were used to construct a minimum spanning tree (MST) and infer lineage structure in an unsupervised manner via fitting of simultaneous principal curves through the MST edges. No start or end clusters were specified. Cell-level pseudotime values were extracted for each inferred lineage. Both continuous pseudotime values and lineage weights were retained for downstream modeling and visualization.

#### Pseudotime alignment and pseudo-bulk lineage visualizations

Pseudotime alignment of the three inferred lineages was performed using dynamic time warping (DTW). Expression of each molecular target was mean-centered and scaled, and smoothed per-lineage expression was estimated using generalized additive models. Lineages 2 and 3 were sequentially aligned to Lineage 1 as reference, first generating a consensus pseudotime between Lineage 1 and 2, that was then used to align Lineage 3. Bifurcations or warped paths across lineages were estimated using pairwise cosine dissimilarity in the shared pseudotime space. For visualization, mean expression was calculated across 50 equally-spaced pseudotime bins per lineage and projected into two dimensions via multidimensional scaling. Differentially expressed molecules were identified using generalized linear models with Tukey tests per bin, and statistically significant results for each molecular target were aggregated across bins using Fisher’s method.

#### Integration of multi-modal single-cell datasets and latent representation learning

Single-nucleus RNA sequencing (snRNA-seq) and single-nucleus ATAC sequencing (snATAC-seq) datasets from E3-AD and E4-AD individuals were integrated to enable joint modeling of transcriptional and chromatin accessibility profiles. The snRNA-seq expression exported from Seurat and the snATAC-seq cell-by-peak count matrices were independently formatted as AnnData objects. Cell barcodes were intersected and harmonized across modalities to define a shared set of matched cells. Both modalities were combined into a single MuData container, with a unified observation table.

Joint probabilistic modeling of the integrated dataset was performed using the MultiVI^86^ framework implemented in scvi-tools (v1.3.3). Donor identity was incorporated as a batch covariate to account for inter-individual variation. The model was configured to learn a 20-dimensional latent representation and trained for 200 epochs with a batch size of 128, yielding a shared RNA-ATAC latent space with donor-corrected structure. The learned MultiVI latent representations of the paired snATAC-seq and snRNA-seq profiles were used to construct a k-nearest neighbor (kNN) graph for uniform manifold approximation and projection (UMAP) visualization and MELD likelihood estimation^47^. For each cell, a difference score (MELD_Δ_ = MELD_4/4_ − MELD_3/3_) was calculated to represent the direction and magnitude of enrichment along the genotype-associated continuum.

Integration performance and biological signal retention were assessed using Local Inverse Simpson’s Index (LISI) across multiple neighborhood sizes. iLISI indicated effective donor mixing, while cLISI demonstrated preservation of APOE-associated structure. Silhouette analysis further supported these results, with reduced donor-driven clustering and modest retention of genotype-specific separation.

#### Assignment of microglia to MELD_Δ_ bins

Cells were stratified based on differential MELD enrichment by applying Gaussian mixture modeling (GMM) to MELD_Δ_ values. A three-component model was used to partition cells into MELD-low, intermediate, and MELD-high groups, corresponding to progressively increasing association with APOE4. This probabilistic clustering approach enabled data-driven discretization of the APOE-association landscape for downstream comparative analyses. The GMM framework was implemented in the *mclust* R package (v6.0.1).

#### Donor-aware differential gene expression analysis

Differential gene expression between MELD-high and MELD-low cells was assessed using a donor-aware pseudobulk approach. Prior to analysis, donors lacking representation in both MELD-high and MELD-low bins were excluded to ensure all donors contributed to both conditions and to prevent confounding of the MELD bin contrast with donor identity. Specifically, one donor (BANN0721) was removed as it contributed cells exclusively to the MELD-high bin, rendering its inclusion in a paired donor-blocking design inappropriate. Raw counts were aggregated across cells for each donor and MELD_Δ_ bin to generate donor-level pseudobulk expression profiles. Low-expression genes were removed prior to statistical testing using the filterByExpr function from *edgeR* (v 4.0.16), which retained genes with sufficient counts across the minimum number of samples required by the experimental design. This filtering step was applied after construction of the design matrix to ensure threshold calculations reflected the actual comparisons being made. Gene-level counts were analyzed using the limma-voom framework (v 3.58.1) with trimmed mean of M values (TMM) normalization implemented in *edgeR* (v 4.0.16). To account for heterogeneity in pseudobulk library sizes arising from unequal cell numbers per donor-bin combination, precision weights were estimated using voomWithQualityWeights, which additionally downweights samples with poor precision relative to the mean-variance trend. Gene-wise linear models were fitted with donor identity included as a blocking factor to account for inter-individual variability. The coefficient corresponding to MELD-high versus MELD-low was extracted to estimate log_2_ fold changes. Statistical significance was determined using empirical Bayes-moderated statistics, and p-values were adjusted using the Benjamini-Hochberg false discovery rate (FDR) procedure.Genes with FDR < 0.05 and absolute log_2_ fold change ≥ 0.5 were considered differentially expressed.

#### Continuous association of gene expression along the APOE continuum

To identify features associated with progression along the APOE genotype axis, gene expression was modeled as a function of MELD_Δ_ and donor identity using a linear modeling framework implemented in the *limma* R package (v 3.58.1). The MELD_Δ_ β coefficient represents the change in log-normalized gene expression associated with a one-unit increase in MELD_Δ_, after adjusting for donor effects. Positive coefficients indicate genes whose expression increases toward the APOE4-associated side of the microglial manifold, while negative coefficients indicate genes enriched toward the APOE3-associated side. Models were fit simultaneously across all genes using lmFit, and empirical Bayes variance moderation was applied using eBayes to improve variance estimation across genes. P-values were adjusted for multiple testing using the Benjamini–Hochberg procedure to control the false discovery rate (FDR). Genes positively associated with MELD_Δ_ were defined using thresholds of FDR < 0.05 and log_2_ fold change ≥ 0.2 and interpreted as increasing toward the APOE4-associated end of the continuum. These genes were then compared against previously defined microglial transcriptional signatures, including Terminally Inflammatory Microglia (TIM)^17^ and homeostatic, DIM, inflammatory DAM, Lipo.DAM, Ribo.DAM1, and Ribo.DAM2 signatures from the Human Microglia Atlas^103^. Overlapping genes were annotated according to their membership in one or more reference signatures. Genes were then ranked by their moderated t-statistic for the MELD_Δ_ coefficient and used as input for downstream gene set enrichment analysis (GSEA).

#### Gene set enrichment analysis (GSEA)

Pre-ranked gene set enrichment analysis was conducted using the *fgsea* package in R^129^ (v1.28.0). Gene sets were retrieved from the *msigdbr* database (v25.1.0) for Homo sapiens, including Reactome (C2:CP:REACTOME), Gene Ontology Biological Process (C5:BP), KEGG Medicus (C2:CP:KEGG_MEDICUS), and Hallmark (H) collections. Each gene set category was converted into a list of pathways keyed by gene set name and containing member gene symbols, and all pathway lists were merged for downstream analysis. Enrichment testing was performed using fgseaMultilevel with the combined pathway collection. Pathways with adjusted p-value < 0.05 and a minimum gene set size of 5 were considered statistically significant. Enrichment direction was determined based on the normalized enrichment score (NES), with positive NES values indicating enrichment toward the APOE4-associated side of the microglial manifold and negative NES values indicating enrichment toward the APOE3-associated side. To identify modules of biologically coherent pathways, we assessed pairwise similarity among the top 50 statistically significant pathways in both enrichment directions. For each pathway, the leading-edge gene set was extracted from the GSEA output. Genes appearing in the leading edge of fewer than two pathways were excluded to focus the analysis on co-regulated pathway modules. Pairwise Jaccard similarity between pathways was computed from a sparse binary pathway-by-gene matrix via matrix cross-product. Pathways were clustered by hierarchical clustering (average linkage) on the resulting distance matrix (1 − Jaccard similarity) and grouped into modules by dendrogram cutting.

#### Differential chromatin accessibility and motif analysis

To derive condition-specific chromatin landscapes, pseudobulk replicates were generated within each MELD group using addGroupCoverages, and peaks were re-called de novo using MACS3 via addReproduciblePeakSet, grouping cells by MELD bin. This produced group-specific peak sets reflecting the distinct open chromatin profiles of MELD-high and MELD-low microglia. Differential chromatin accessibility between MELD-high and MELD-low cells was assessed using getMarkerFeatures on the PeakMatrix. To assess differences in transcription factor (TF) motif accessibility, motif enrichment analysis was performed using chromVAR^95^ with motif annotations derived from the CIS-BP database. GC-matched background peaks were selected using addBgdPeaks, and per-cell TF motif deviation scores were computed with addDeviationMatrix, generating a MotifMatrix within the ArchR project. Differential TF motif deviation between MELD-high and MELD-low groups was then assessed using getMarkerFeatures on the MotifMatrix, applying significance thresholds of FDR ≤ 0.05 and mean difference ≥ 0.03.

## DATA AVAILABILITY

All data generated in this work will be made publicly available upon manuscript acceptance.

