## Supplemental Data and Tables for "APOE4 Drives Uniquely Dysfunctional Human Microglial States in Alzheimer’s Disease"

#### Supplementary Figure 1

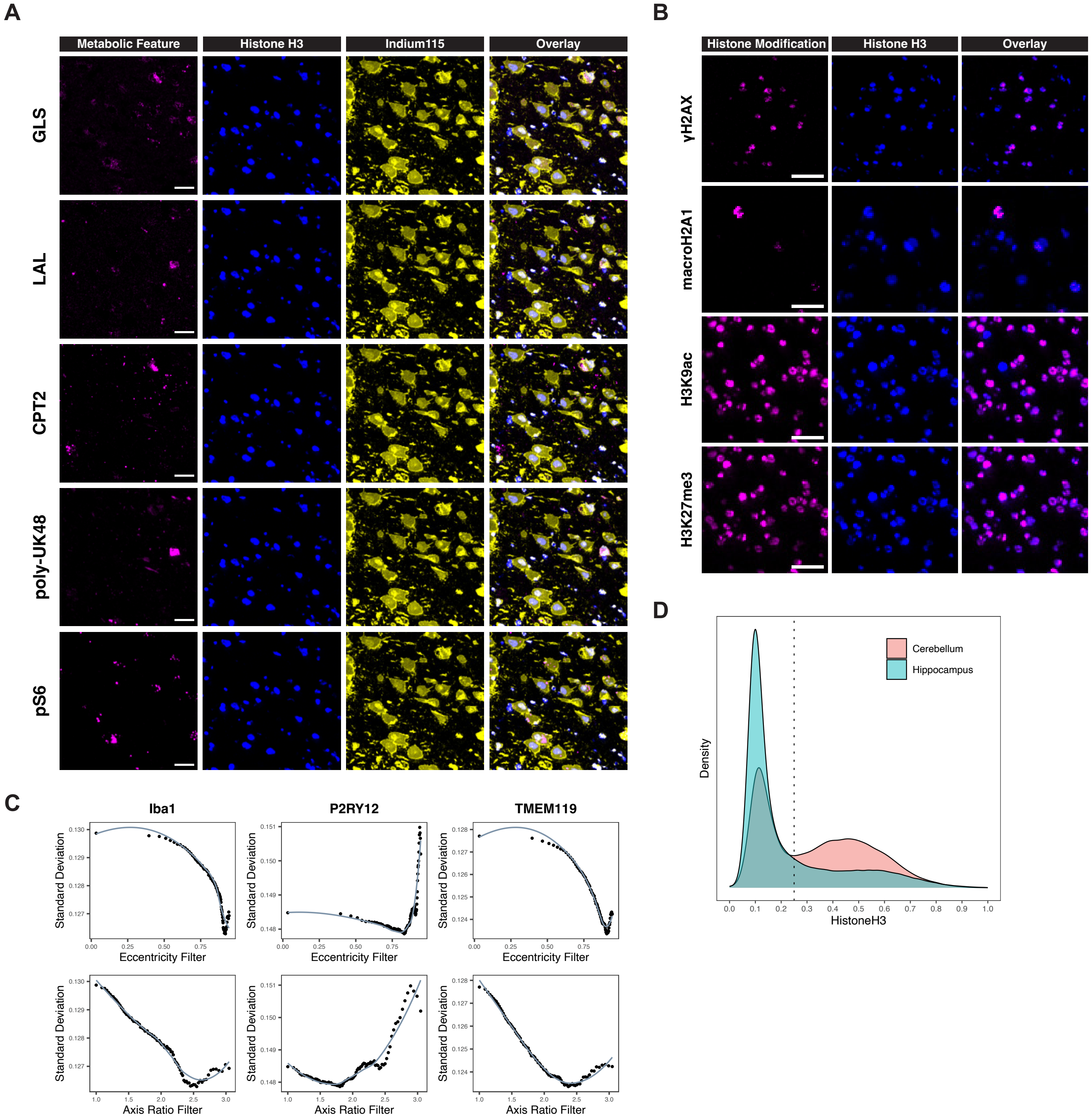


**(A)** Representative MIBI image demonstrating cytoplasmic localization of metabolic proteins within pyramidal neurons. Scale bars in multi-panel single-cell callout images (bottom) represent 10um. **(B)** Representative MIBI image demonstrating nuclear localization of histone modification markers. Scale bars in multi-panel single-cell callout images (bottom) represent 10um. **(C)** To refine morphological gating of anucleated microglia and exclude small transverse sections of glial processes, we systematically applied percentile-based morpho-filters across eccentricity and major-to-minor axis ratio. Specifically, for each percentile threshold n in the range 1-99, cells below the nth percentile for each parameter were progressively excluded. At each filtering step, we quantified the standard deviation (SD) of canonical microglial markers (Iba1, P2RY12, TMEM119) to assess marker stability under increasing stringency. The percentile cutoff at which the SD of marker expression reached a global minimum was identified as the optimal morphological filter. Based on these results, we retained only anucleated microglia that satisfied the following criteria: eccentricity ≥ 0.85 and major-to-minor axis ratio ≥ 2. **(D)** Density distribution of Histone H3 expression of segmented cellular objects across cerebellum and hippocampal tissue samples. Dotted line indicates cutoff value of 0.25 for defining nucleated cellular objects.

#### Supplementary Figure 2

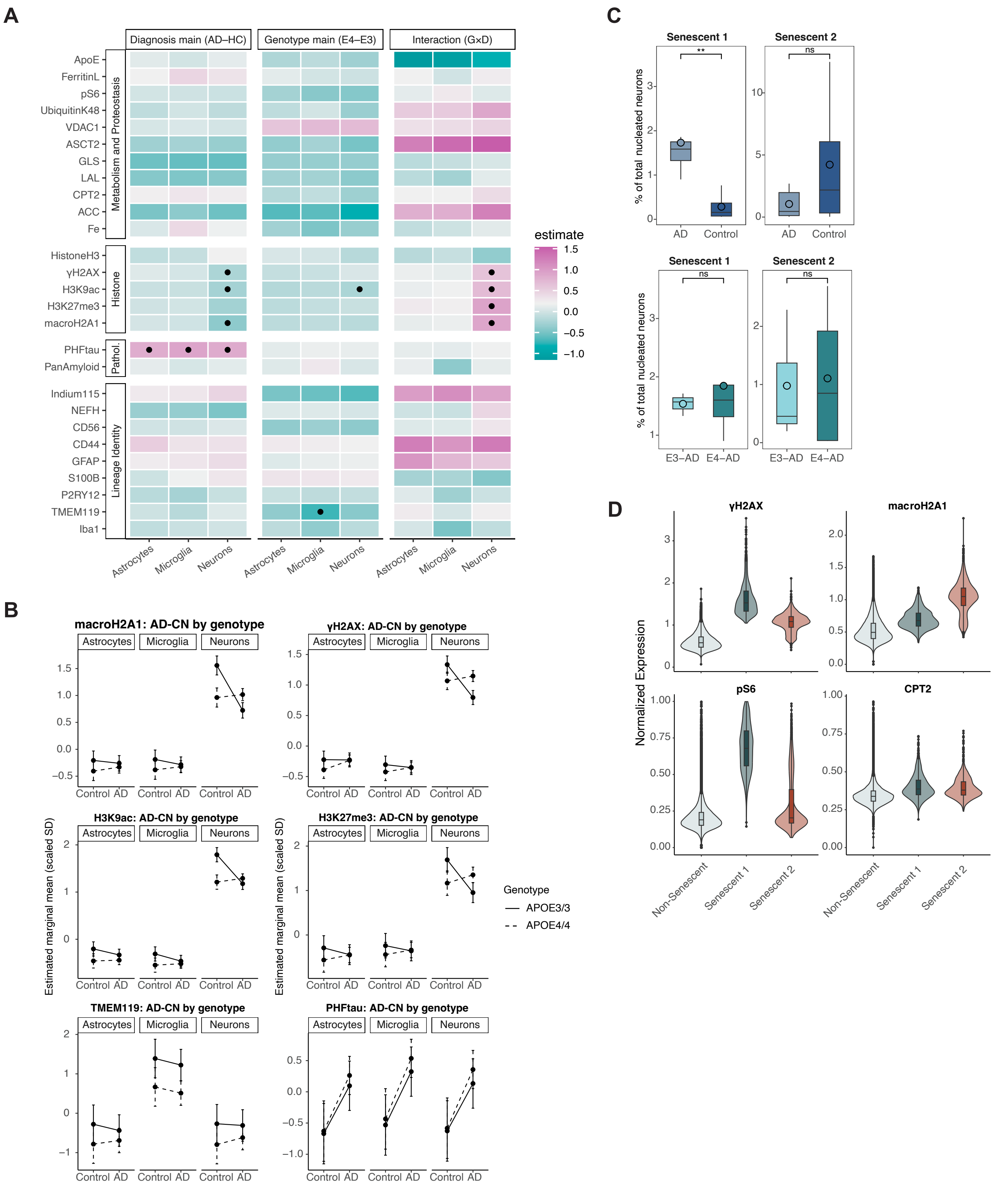

**(A)** Signed effect-size heatmaps. Tiles display signed estimates (SD units) for the genotype main effect (E4-E3, averaged over diagnosis), the diagnosis main effect (AD-CN, averaged over genotype), and the G×D interaction defined as (E4,AD-E4,CN)-(E3,AD-E3,CN). Dots denote effects meeting both statistical significance (FDR < 0.05) and the 0.2 SD relevance threshold. **(B)** Interaction plot of estimated marginal means from the linear mixed-effects model. Line type denotes genotype; points are model-based means with 95% confidence intervals. Non-parallel lines indicate a genotype-diagnosis interaction; vertical separation between the line types at a given diagnosis reflects the genotype main effect; overall vertical shift from CN to AD reflects the diagnosis main effect. **(C)** Senescent 1 was found to be significantly enriched in AD neurons, implicating this population in disease-associated senescence, although there was no significant difference in abundance between E3-AD and E4-AD. **(D)** Senescent 1 was found to show elevated levels of γH2A.X and reduced macroH2A1 positivity relative to Senescent 2.

#### Supplementary Figure 3

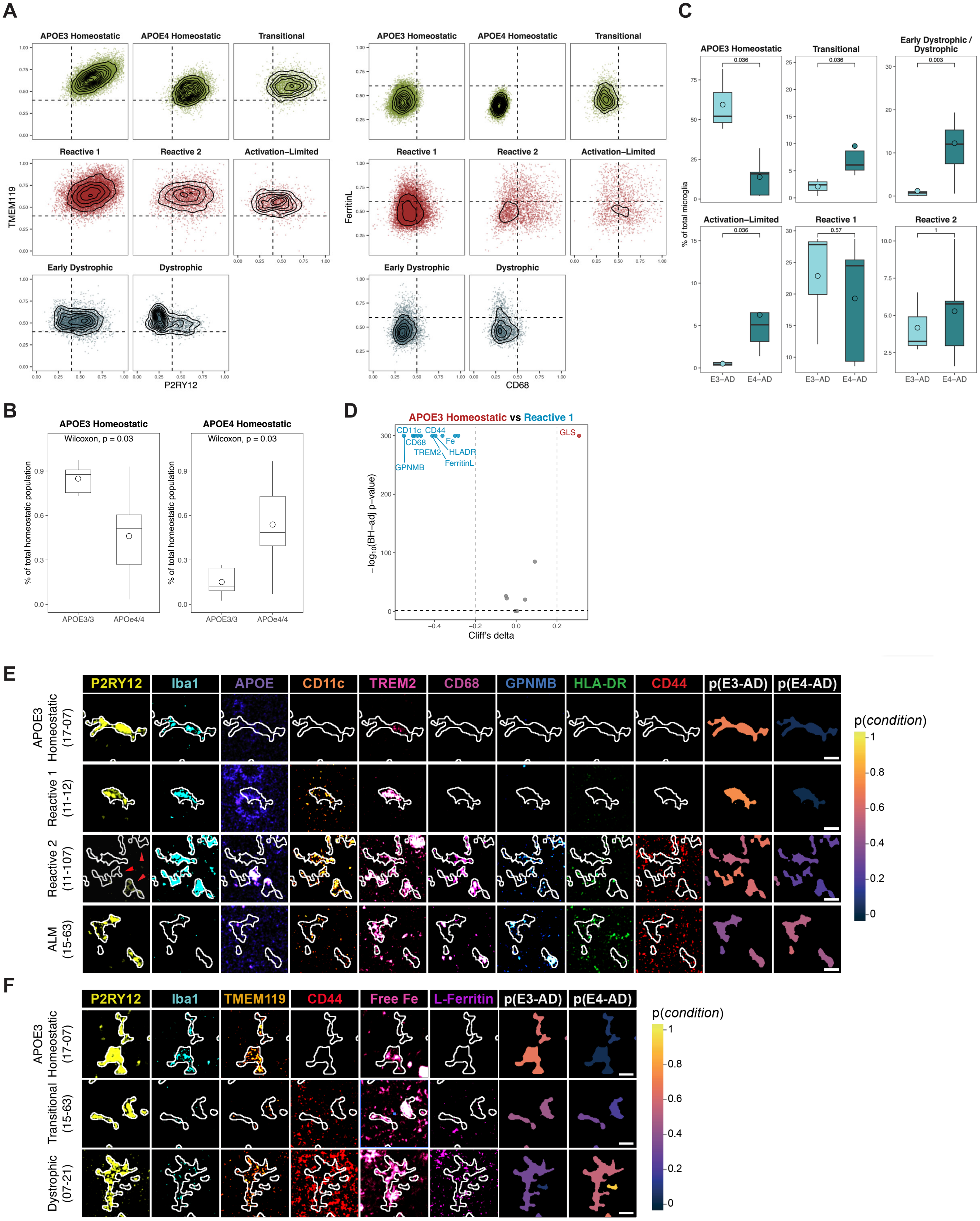


**(A)** Biaxial scatter plots of TMEM119 vs P2RY12, faceted by cluster (left). Clusters in the non-activated group (top row) show high expression of both markers. Reactive clusters (middle row) show partial reduction, and dystrophic clusters (bottom row) exhibit pronounced loss, supporting the phenotypic group classification. Biaxial scatter plots of L-ferritin vs CD68, faceted by cluster (right). Clusters in the non-activated group show low expression of both markers, reactive clusters show elevated CD68 with emerging L-ferritin, and dystrophic clusters exhibit high L-ferritin, supporting the phenotypic group classification. **(B)** Frequencies of APOE3- vs APOE4-associated homeostatic clusters per individual, expressed as a percentage of the total homeostatic population. APOE3 Homeostatic cells are enriched in APOE3/3 individuals, whereas APOE4 Homeostatic cells are enriched in APOE4/4 individuals (Wilcoxon Rank-Sum $p < 0.05$). **(C)** Boxplots of cluster frequencies show that while Reactive 1 and Reactive 2 do not differ between genotypes, ALMs are specifically enriched in E4-AD individuals. E4-AD individuals show depletion of canonical homeostatic states in concert with an increase in APOE4-specific transitional and dystrophic states. **(D)** Volcano plot showing differential expression of key microglial activation markers between APOE3 Homeostatic and Reactive 1. **(E)** Representative multiplexed images of Reactive 1, Reactive 2 and ALMs. **(F)** Representative multiplexed images of Transitional and Dystrophic cells.

### Supplementary Figure 4

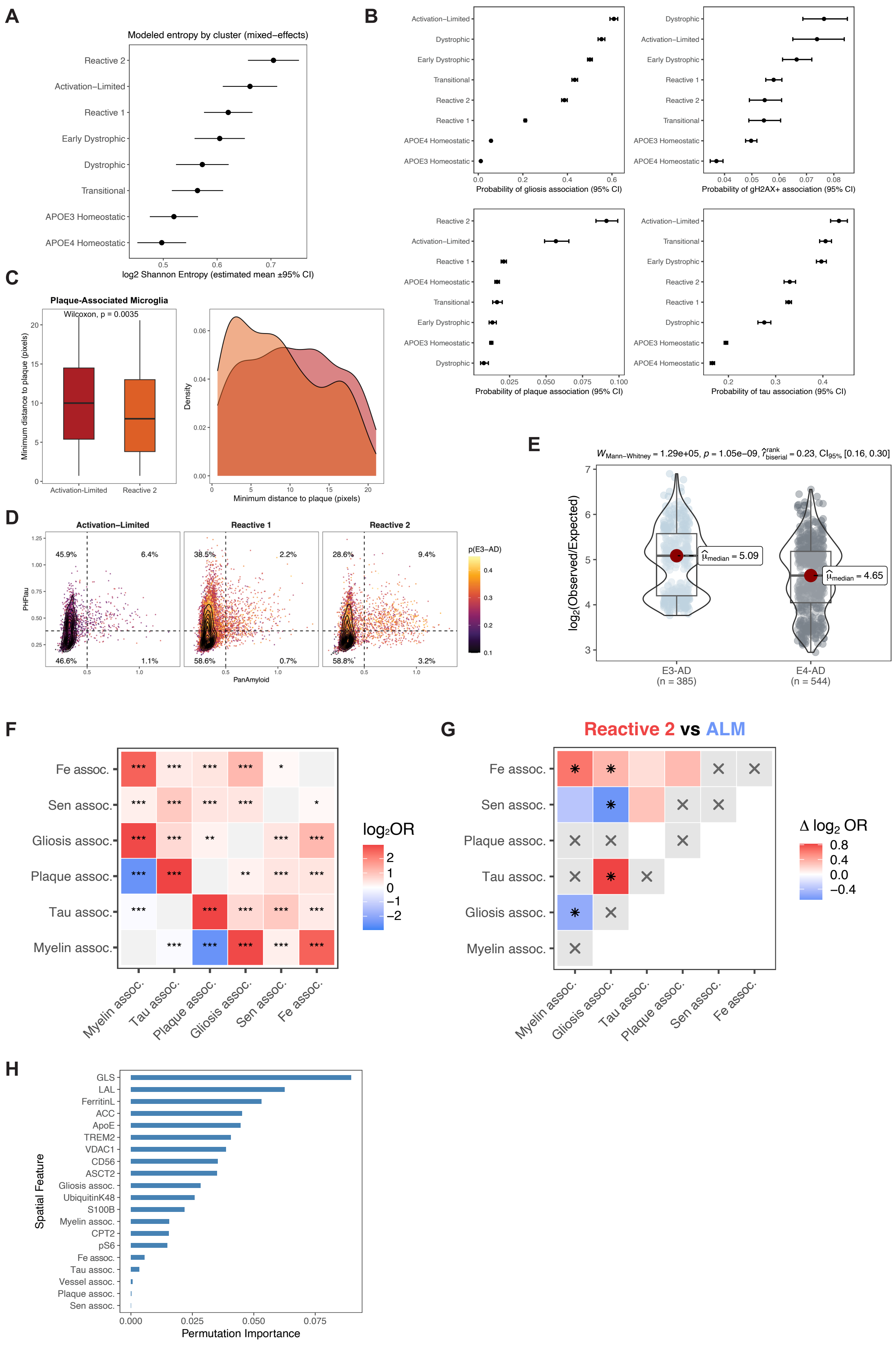

**(A)** $Log_2$ Shannon entropy per cell was modeled using a linear mixed-effects model with cluster as a fixed effect and donor as a random intercept. Points show estimated marginal means (±95% CI), adjusted for sample-level random effects. Clusters are ordered by estimated mean entropy. **(B)** Logistic regression odds ratios with 95% confidence intervals for spatial feature association by cluster. ALMs were more likely to be tau-associated relative to Reactive 2 microglia, whereas Reactive 2 microglia were significantly enriched for plaque association. Senescent (gH2AX+) neighbors were strongly associated with dystrophic and ALM states, indicating cluster-specific spatial pathology. **(C)** Spatial distance-to-plaque distributions. Reactive 2 microglia are significantly closer to amyloid plaques, consistent with plaque-associated DAM identity; ALMs remain farther away, reflecting non-plaque localization. **(D)** Spatial neighborhood analysis of DAMs versus ALMs. Points represent the mean pixel expression profile in a custom 20-pixel expansion area around individual microglia. Reactive 2 microglia preferentially localize near amyloid plaques (12.6%), consistent with plaque-associated DAM identity and a role in focal neuroinflammation. On the other hand, only 7.5% of ALMs are plaque-associated, underscoring divergent microenvironmental contexts. **(E)** Microglia in E3-AD are more spatially concentrated around plaques than in E4-AD. To quantify microglial coverage around plaques, we calculated the expected area of microglia in the expansion region assuming uniform distribution across the field of view (FOV). We then compared this to the observed microglial area within each plaque's 50-pixel expansion zone. The resulting enrichment ratio – defined as the log2 ratio of observed to expected microglial area coverage – served as a measure of microglial recruitment to plaques. Positive values indicate local enrichment, while negative values reflect under-representative relative to a uniform spatial distribution. This metric allows for direct comparison across clusters or plaque types while accounting for baseline microglial density in the tissue. **(F)** Global co-occurrence heatmap of spatial features. Each cell shows the log2 odds ratio for pairwise co-occurrence across all microglia (Fisher's exact test); positive values indicate enrichment; negative values indicate mutual exclusivity. Non-significant pairs (FDR ≥ 0.05) are muted. **(G)** Heatmap of differential co-occurrence between Reactive 2 and ALM clusters, showing $\Delta log_2 OR$ for each spatial feature pair. A 2×2×2 log-linear model tested whether the strength of association between each feature pair differed by genotype. Stars denote FDR-adjusted significance. **(H)** Feature permutation importance scores from a random forest classifier predicting microglial cluster identity from spatial features. Among the top-ranked predictors were pixel-level metabolic markers (e.g., GLS, LAL, APOE, VDAC1, TREM2) extracted from the neighborhood expansion region, underscoring the contribution of local metabolic tone to microglial state.

#### Supplementary Figure 5

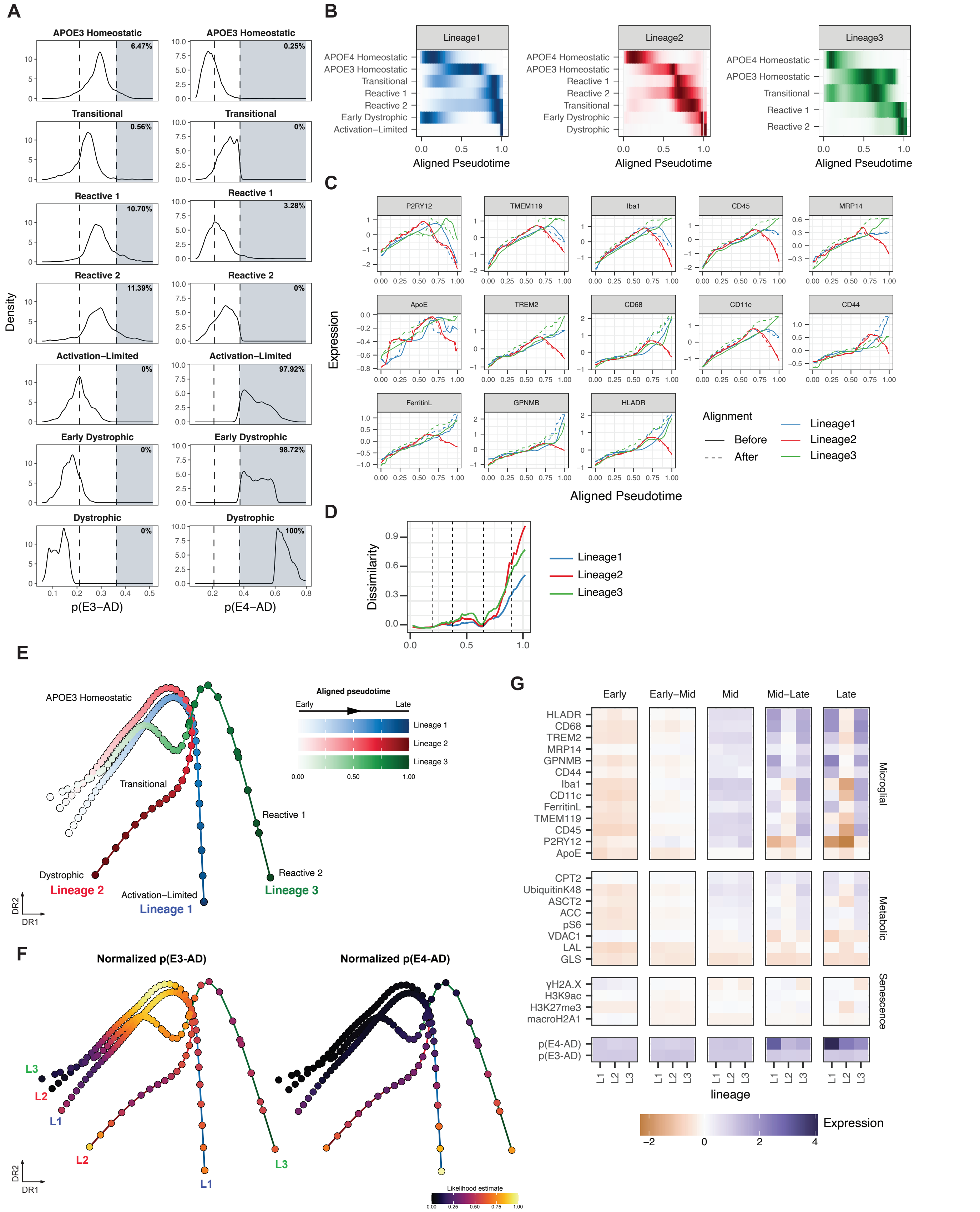


**(A)** Density distribution along the *p*(E3-AD) and *p*(E4-AD) continua; dotted lines denote the boundaries for low, middle, and high likelihood bins. Thresholds were determined via Gaussian Mixture Modeling of each likelihood continuum with k = 3 components. Percentage of cells per cluster found in the high likelihood bin is denoted in the top right corner or each plot. Likelihood distributions reveal a graded shift along the *p*(E4-AD) continuum: homeostatic clusters are confined to low likelihoods, Reactive 1 and Reactive 2 occupy intermediate ranges, and ALMs dominate the far-right tail, reflecting their strong and distinct alignment with E4-AD. Homeostatic cells localize to low likelihoods, while Transitional and Dystrophic states are increasingly enriched at high *p*(E4-AD). **(B)** Distribution of cellular states across aligned pseudotime in the three inferred lineages. Rows correspond to annotated cell clusters (as applicable per lineage). Color intensity reflects the relative enrichment (density) of cells from each cluster at a given pseudotime point within each lineage, with darker shades indicating higher density. **(C)** Comparison of marker expression across pseudotime for each lineage prior to- (solid line) and post-trajectory alignment (dotted line). **(D)** Mean cosine dissimilarity between feature expression profiles, computed for each lineage relative to the other two. Vertical dotted lines indicate manually defined pseudotime bins. **(E)** Lineage embeddings constructed from the mean expression of microglial features within pseudotime bins along each aligned trajectory. All three lineages share a similar early topology, before diverging into distinct terminal phenotypes. **(F)** Lineage embeddings colored by normalized MELD condition-associated likelihood scores for E3-AD (left) and E4-AD (right). Select pseudotime bins are labeled by their predominant cluster identity. **(G)** Expression heatmap of features across aligned pseudotime for all lineages, stratified by pseudotime bin.

#### Supplementary Figure 6

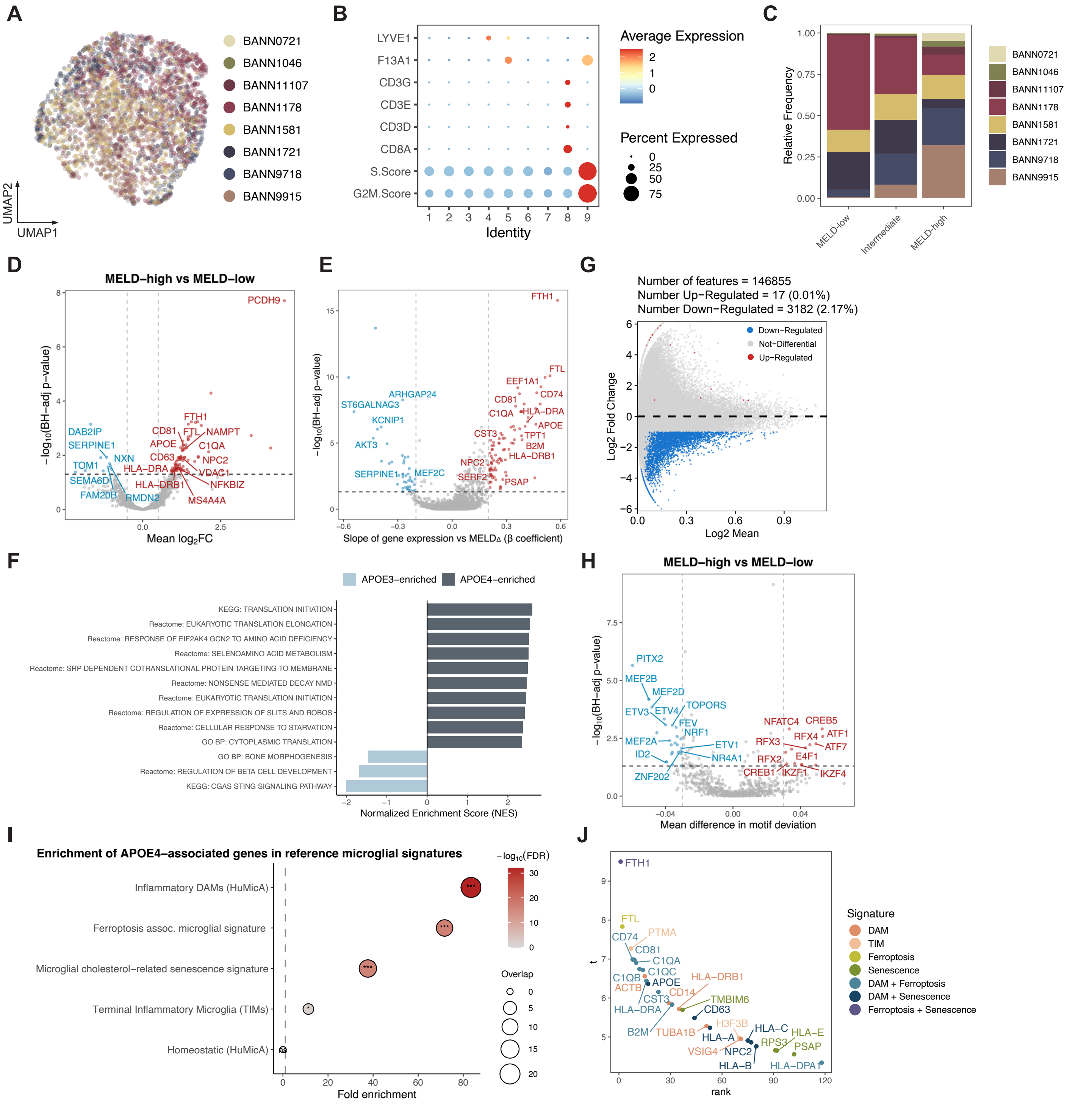


**(A)** UMAP colored by donor, demonstrating donor mixing following MultiVI integration. **(B)** Identification of cycling microglia and T cell populations. The balloon plot of Leiden clusters shows cluster 9 enriched for cycling cells (high G2M and S scores) and cluster 8 enriched for T cell markers (CD3D-G, CD8A). Clusters 8 and 9 were removed prior to MultiVI integration. **(C)** Donor contribution across $\text{MELD}_\Delta$ bins. **(D)** Differential gene expression between MELD-high and MELD-low cells. Pseudobulk counts were aggregated by donor and $\text{MELD}_\Delta$ bin prior to analysis with a limma-voom framework. Differentially expressed genes (FDR < 0.05, $\log_2$FC ≥ 0.5) are highlighted in red. **(E)** Continuous association analysis along $\text{MELD}_\Delta$. Each point represents a gene; coefficients indicate direction of association with the APOE continuum (positive: MELD-high; negative: MELD-low). Significant genes (FDR < 0.05 and |β| ≥ 0.2) are highlighted. **(F)** Genes were ranked by their moderated tstatistics from donor-aware continuous $MELD_{\Delta}$ association analysis. The normalized enrichment score (NES) plot shows top 10 pathways enriched at each end of the continuum. Positive NES values indicate pathways upregulated in MELD-high (APOE4-associated) cells, while negative NES values indicate pathways enriched in MELD-low (APOE3-associated) cells. Significance was assessed using FDR-adjusted p-values. **(G)** Pairwise differential peak accessibility between MELD-high and MELD-low groups shown as an MA plot, illustrating effect size versus mean accessibility. **(H)** Differential transcription factor (TF) motif accessibility between MELD-high and MELD-low cells accessed using chromVAR motif deviation scores. The volcano plot shows mean deviation differences versus −log10(FDR), with significantly different motifs (FDR < 0.05, mean difference ≥ 0.03) highlighted in red. **(I)** Bubble plot showing enrichment of the APOE4-associated gene set across published microglial reference signatures. The x-axis indicates fold enrichment relative to expectation under the background gene universe, point size denotes the number of overlapping genes, and color denotes FDR-adjusted significance from one-sided Fisher's exact tests. **(J)** Genes from the APOE4 signature that overlap with a reference microglial signature are shown as individual points, colored by signature membership. The x-axis indicates gene rank based on GSEA ordering, and the y-axis indicates the moderated t-statistic from continuous association analysis as shown in (E), illustrating the relative positioning of inflammatory DAM, ferroptosis-, and senescence-associated genes along the APOE4-associated transcriptional axis.

**Supplementary Figure 7**

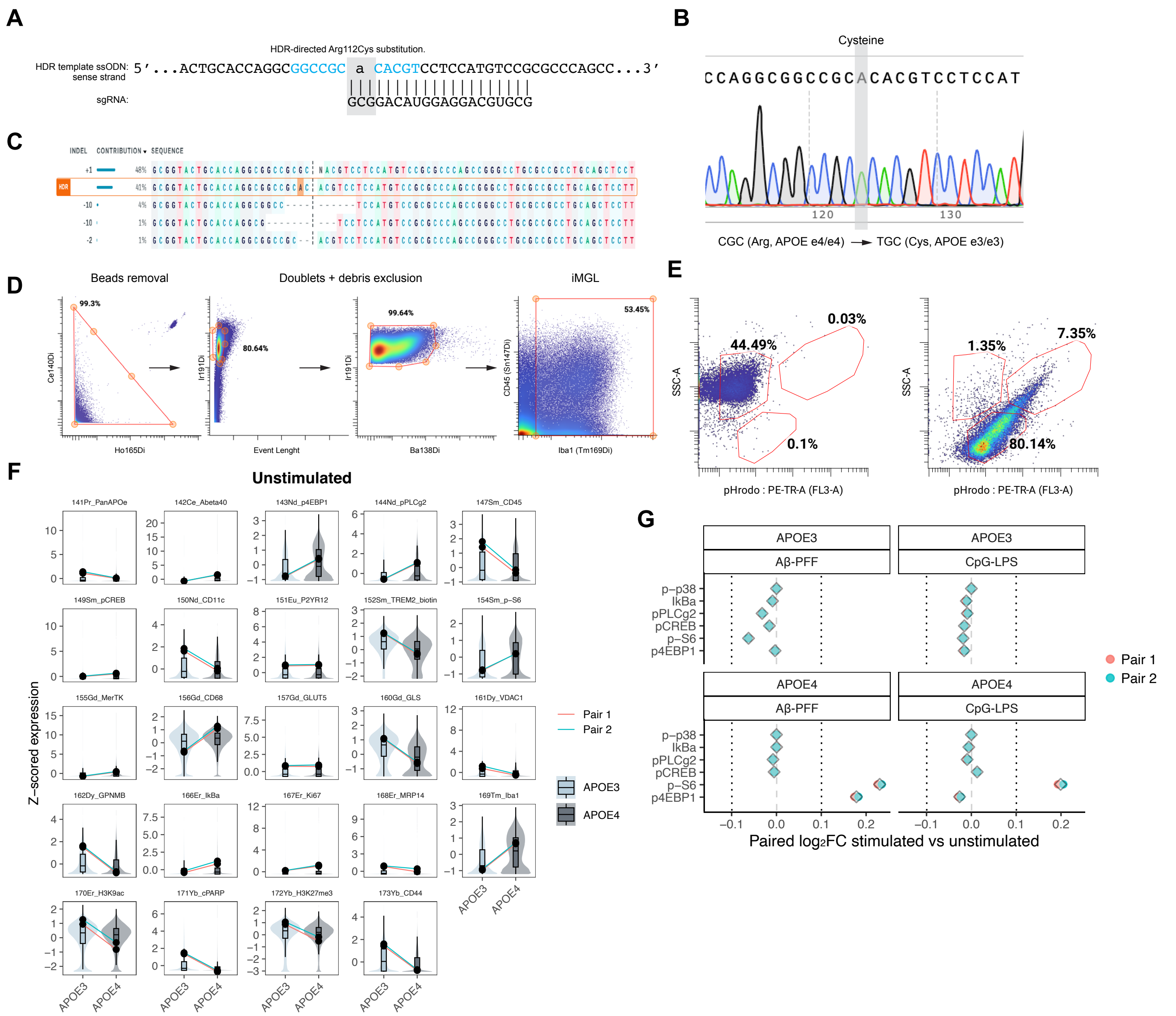


**(A)** Design of CRISPR/Cas9-mediated homology-directed repair (HDR) to introduce the APOE ε3 allele. The ssODN repair template encodes the Arg112Cys substitution (CGC→TGC), with the sgRNA targeting sequence shown aligned to the genomic locus. **(B)** Sanger sequencing chromatogram confirming successful editing at the APOE locus. The nucleotide substitution corresponding to Arg→Cys is highlighted, demonstrating conversion from APOE ε4/ε4 (CGC) to APOE ε3/ε3 (TGC). **(C)** Sequence decomposition analysis of edited populations showing HDR-mediated incorporation alongside indel profiles. The relative contribution of edited and indel-containing alleles is displayed, with the HDR sequence highlighted. **(D)** CyTOF gating workflow for identification of iMGLs. Sequential gating includes bead removal, exclusion of doublets and debris based on event length and DNA content, followed by identification of iMGLs based on CD45 and Iba1 expression. Percentages indicate the proportion of events retained at each step. **(E)** Flow cytometry gating strategy for the iMGL phagocytosis assay. Representative plots show gating of pHrodo-positive cells based on SSC-A versus pHrodo (PE-TR-A) fluorescence, with percentages indicating gated populations. **(F)** Comprehensive marker-level distributions of unstimulated microglia, stratified by APOE genotype. Violin and box plots display single-cell distributions for all measured markers across conditions and genotypes. Overlaid points indicate z-scored pseudobulked medians. Lines connect matched isogenic pairs, emphasizing consistency of observed genotype-driven differences across replicates. **(G)** Stimulus-induced responses by APOE genotype. Paired

$\log_2$ fold-change ($\log_2$FC) in marker expression following stimulation (Aβ/PFF or CpG/LPS, 12 min) relative to matched unstimulated controls, shown separately for APOE3 and APOE4 microglia. Each point represents the paired response for an individual isogenic pair; diamond-shaped points summarize the mean across pairs. Vertical dashed lines indicate no change (0) and reference thresholds (±0.1).

### Supplementary Table 1

| apoe_diag | AD_3 | AD_3 | AD_3 | AD_4 | AD_4 | AD_4 | AD_4 | AD_4 | control_3 | control_3 | control_4 | control_4 |
|---|---|---|---|---|---|---|---|---|---|---|---|---|
| CaseID | 11-12 | 11-78 | 08-20 | 07-21 | 08-71 | 97-18 | 11-107 | 15-63 | 17-50 | 17-07 | 99-02 | 97-09 |
| race | 1 | 1 | 1 | 1 | 1 | 1 | 1 | 1 | 1 | 1 | 1 | 1 |
| gender | 2 | 1 | 1 | 2 | 1 | 2 | 1 | 2 | 1 | 2 | 2 | 2 |
| expired_age | 76 | 82 | 76 | 92 | 85 | 93 | 71 | 80 | 90 | 76 | 70 | 81 |
| PMI | 2 | 2.95 | 3.33 | 2.83 | 13 | 2.5 | 4.32 | 19 | 3.41 | 2.93 | 2 | 3 |
| last_mmse_test_score | | 19 | 16 | 13 | 14 | | 0 | 19 | 27 | 30 | | |
| last_mmse_interval_before_death | | 16 | 13 | 32 | 17 | | 2 | 38 | 54 | 3 | | |
| Control | no | no | no | no | no | no | no | no | yes | yes | yes | yes |
| AD | yes | yes | yes | yes | yes | yes | yes | yes | no | no | no | no |
| PD | no | no | no | no | no | no | no | no | no | no | no | no |
| DLB | no | no | no | no | no | no | no | no | no | no | no | no |
| VAD | no | no | no | no | no | no | no | no | no | no | no | no |
| PSP | no | no | no | no | no | no | no | no | no | no | no | no |
| HS | no | no | no | no | no | no | no | no | no | no | no | no |
| DLDH | no | no | no | no | no | no | no | no | no | no | no | no |
| MND | no | no | no | no | no | no | no | no | no | no | no | no |
| CBD | no | no | no | no | no | no | no | no | no | no | no | no |
| PICKS | no | no | no | no | no | no | no | no | no | no | no | no |
| HD | no | no | no | no | no | no | no | no | no | no | no | no |
| MSA | no | no | no | no | no | no | no | no | no | no | no | no |
| ARG | no | no | no | no | no | no | no | no | no | no | no | no |
| CWMR | yes | yes | yes | yes | no | yes | yes | yes | no | yes | no | no |
| dementia_nos | no | no | no | no | no | no | no | no | no | no | no | no |
| MS | no | no | no | no | no | no | no | no | no | no | no | no |
| CAA | no | yes | yes | yes | yes | yes | yes | yes | no | yes | yes | yes |
| MCI | no | no | no | no | no | no | no | no | no | no | no | no |
| LBS | no | no | no | no | no | no | no | no | no | no | no | no |
| BRAIN_ABNORMAL | yes | no | no | no | no | no | no | no | no | yes | no | no |
| ACUTE_INFARCTS | no | no | no | no | no | no | no | no | no | no | no | no |
| ApoE | 3/3 | 3/3 | 3/3 | 4/4 | 4/4 | 4/4 | 4/4 | 4/4 | 3/3 | 3/3 | 4/4 | 4/4 |
| PlaqueF | 3 | 3 | 3 | 3 | 3 | 3 | 3 | 3 | 0.5 | 0 | 1.5 | 2.5 |
| PlaqueT | 3 | 3 | 3 | 3 | 3 | 3 | 3 | 3 | 0 | 0 | 2 | 1.5 |
| PlaqueP | 3 | 3 | 3 | 3 | 3 | 2.5 | 3 | 3 | 0 | 0 | 2 | 1.5 |
| PlaqueH | 3 | 2.5 | 3 | 2.5 | 3 | 2.5 | 2.5 | 2 | 0 | 0 | 1 | 1 |
| PlaqueE | 3 | 3 | 3 | 3 | 3 | 3 | 2.5 | 3 | 0 | 0 | 1.75 | 1 |
| PlaqueTotal | 15 | 14.5 | 15 | 14.5 | 15 | 14 | 14 | 14 | 0.5 | 0 | 8.25 | 7.5 |
| Plaque density | frequent | frequent | frequent | frequent | frequent | frequent | frequent | frequent | sparse | zero | sparse | sparse |
| TangleF | 3 | 3 | 1.5 | 3 | 3 | 3 | 3 | 3 | 0 | 0 | 0 | 0 |
| TangleT | 3 | 3 | 3 | 3 | 3 | 3 | 3 | 3 | 0 | 0 | 0 | 0 |
| TangleP | 3 | 3 | 2 | 3 | 3 | 2.5 | 3 | 3 | 0 | 0.5 | 0 | 0 |
| TangleE | 3 | 3 | 3 | 3 | 3 | 3 | 3 | 3 | 1 | 1.5 | 0 | 1 |
| TangleH | 3 | 3 | 3 | 3 | 3 | 3 | 3 | 3 | 0.5 | 1.5 | 0 | 2 |
| TangleTotal | 15 | 15 | 12.5 | 15 | 15 | 14.5 | 15 | 15 | 1.5 | 3.5 | 0 | 3 |
| Cerad NP | definite AD | definite AD | definite AD | definite AD | definite AD | definite AD | definite AD | definite AD | Criteria not met | Criteria not met | not AD | not AD |
| Braak score | VI | V | V | VI | VI | VI | VI | VI | I | III | I | II |
| NIA-R | high | high | high | high | high | high | high | high | Criteria not met | Criteria not met | criteria not met | criteria not met |
| Unified LB Stage | 0. No Lewy bodies | 0. No Lewy bodies | 0. No Lewy bodies | 0. No Lewy bodies | 0. No Lewy bodies | | 0. No Lewy bodies | 0. No Lewy bodies | 0. No Lewy bodies | 0. No Lewy bodies | 0. No Lewy bodies | 0. No Lewy bodies |
| infarct_total_volume | 0 | 0 | 0 | 1 | 0 | 6.4 | 0 | 0 | 0.1 | 0 | 0 | 0 |
| infarct_cerebral_total_volume | 0 | 0 | 0 | 1 | 0 | 5.7 | 0 | 0 | 0.1 | 0 | 0 | 0 |
| obt | 0 | 0 | 0 | 0 | 0 | | 0 | 0 | 0 | 0 | 0 | 0 |
| brain_stem_ix_x | 0 | 0 | 0 | 0 | 0 | | 0 | 0 | 0 | 0 | | 0 |
| brain_stem_lc | 0 | 0 | 0 | 0 | 0 | | 0 | 0 | 0 | 0 | 0 | 0 |
| bf_amygdala | 0 | 0 | 0 | 0 | 0 | | 0 | 0 | 0 | 0 | 0 | 0 |
| bf_nbm | | | 0 | | | | | | | | | |
| brain_stem_sn | 0 | 0 | 0 | 0 | 0 | | 0 | 0 | 0 | 0 | 0 | 0 |
| bf_trans | 0 | 0 | 0 | 0 | 0 | | 0 | 0 | 0 | 0 | 0 | 0 |
| bf_cing | 0 | 0 | 0 | 0 | 0 | | 0 | 0 | 0 | 0 | 0 | 0 |
| nctx_temporal | 0 | 0 | 0 | 0 | 0 | | 0 | 0 | 0 | 0 | 0 | 0 |
| nctx_frontal | 0 | 0 | 0 | 0 | 0 | | 0 | 0 | 0 | 0 | 0 | 0 |
| sum_lb_density | 0 | 0 | 0 | 0 | 0 | | 0 | 0 | 0 | 0 | | 0 |
| nctx_parietal | 0 | 0 | 0 | 0 | 0 | | 0 | 0 | 0 | 0 | 0 | 0 |
| PathDXSummary | Alzheimer's Disease; Cerebral white matter rarefaction | Alzheimer's disease; Cerebral white matter rarefaction | Alzheimer's disease | Alzheimer's disease; Old lacunar infarct, left putamen; Cerebral white matter rarefaction | Alzheimer's disease with severe amyloid angiopathy; Marked loss of Purkinje cells, cerebellar cortex | Alzheimer's disease; multiple, small old infarcts of cerebral and cerebellar cortex | Alzheimer's disease; Cerebral white matter rarefaction | Alzheimer's disease; Cerebral white matter rarefaction | Control; Microscopic changes of Alzheimer's disease, insufficient for diagnosis; Old microscopic infarct, left middle frontal gyrus | Extensive leptomeningeal carcinomatosis, brain and spinal cord (history of invasive ductal carcinoma, right breast); Cerebral white matter rarefaction; Neurofibrillary tangles, mesial temporal lobe; Amyloid angiopathy, | Control; Alzheimer type II astrocytosis, consistent with terminal hepatic failure | Control |
| adcc_status | NA | NA | NA | NA | Accepted | NA | NA | Accepted | NA | NA | NA | NA |
| neurological_dx | Alzheimer's disease | Alzheimer's disease | Alzheimer's disease | Alzheimer's disease | Alzheimer's disease | Alzheimer's disease | Alzheimer's disease | Alzheimer's disease | Control | Control | Control | Control |
| neurological_dx_years | 15 | 8 | 3 | 8 | 6 | 3 | 10 | 11 | | | | |
| neurological_dx_age | 61 | 74 | 73 | 84 | 79 | 90 | 61 | 69 | | | | |
| dementia_yn | 1 | 1 | 1 | 1 | 1 | 1 | 1 | 1 | 0 | 0 | 0 | 0 |
| dementia_years | 15 | 8 | 3 | 8 | 6 | 3 | 10 | 4 | | | | |
| demential_age | 61 | 74 | 73 | 84 | 79 | 90 | 61 | 76 | | | | |

#### Supplementary Table 2

| Antigen | Clone | Catalog | Source | Isotope | Element | Concentration (ug/ml) |
|---|---|---|---|---|---|---|
| 8OHG | 15A3 | ab62623 | Abcam | 69 | Ga | 1 |
| HH3 | D1H2 | 4499BF | Cell Signaling Technology | 89 | Y | 2 |
| CD56 | MRQ42 | 156R-9-OEM | Cell Marque | 113 | In | 2 |
| Pan APOE | D7I9N | 13366BF | Cell Signaling Technology | 141 | Pr | 1 |
| Abeta40 | BDI350 | ab20068 | Abcam | 142 | Nd | 1 |
| ASCT2 | CAL33 | ab237704 | Abcam | 143 | Nd | 1 |
| ACC | C83B10 | 52923SF | Cell Signaling Technology | 144 | Nd | 1 |
| CPT2 | EPR13626 | ab231162 | Abcam | 145 | Nd | 1 |
| Abeta42 | 12F4 | 805501 | Biolegend | 146 | Nd | 1 |
| CD45 | D9M8I | 47937SF | Cell Signaling Technology | 147 | Sm | 1 |
| CD105 | Poly | AF1097 | R&D Systems | 148 | Nd | 1 |
| CD31 | EP3095 | ab226157 | Abcam | 148 | Nd | 1 |
| GLS | EP7212 | ab214802 | Abcam | 149 | Sm | 1 |
| polyUbiquitin48 | EP8589 | ab221211 | Abcam | 150 | Nd | 0.5 |
| P2RY12 | Poly | NBP2-33870 | Novus | 151 | Eu | 2 |
| Biotin* | 1D4-C5 | 409002 | Biolegend | 152 | Sm | 1 |
| CD11c | EP1347Y | ab216655 | Abcam | 153 | Eu | 1 |
| p-S6 | D57.2.2E | 4858BF | Cell Signaling Technology | 154 | Sm | 0.5 |
| MerTK | Y323 | ab176887 | Abcam | 155 | Gd | 1 |
| CD68 | D4B9C | 76437BF | Cell Signaling Technology | 156 | Gd | 0.5 |
| GLUT5 | E-2 | sc-271055 | Santa Cruz Bio | 157 | Gd | 0.5 |
| gH2A.X | Poly | NB100-384 | Novus | 158 | Gd | 1 |
| NEFH | RMdO 20 | 2836BF | Cell Signaling Technology | 159 | Tb | 0.5 |
| macroH2A1.2 | 14G7 | MABE61 | Millipore Sigma | 160 | Gd | 1 |
| VDAC1 | 20B12AF2 | ab14734 | Abcam | 161 | Dy | 1 |
| GPNMB | Poly | AF2550 | R&D Systems | 162 | Dy | 1 |
| L-ferritin | D-9 | sc-74513 | Santa Cruz Bio | 163 | Dy | 1 |
| TMEM119 | CL8714 | 41134BF | Cell Signaling Technology | 164 | Dy | 1 |
| MCNPase | SMI91 | 836404 | Biolegend | 165 | Ho | 2 |
| LAL | 9G7F12 | ab36597 | Abcam | 166 | Er | 1 |
| HLA-DR | EPR3692 | ab209968 | Abcam | 167 | Er | 0.5 |
| MRP14 | Poly | LS_B12844 | LS Bio | 168 | Er | 0.125 |
| Iba1 | EPR16588 | ab220815 | Abcam | 169 | Tm | 1 |
| H3K9ac | C5B11 | 9649BF | Cell Signaling Technology | 170 | Er | 1 |
| S100b | E7C3A | 90393BF | Cell Signaling Technology | 171 | Yb | 1 |
| H3K27me3 | C36B11 | 9733BF | Cell Signaling Technology | 172 | Yb | 1 |
| CD44 | IM7 | 103002 | Biolegend | 173 | Yb | 2 |
| GFAP | E4L7M | 80788BF | Cell Signaling Technology | 174 | Yb | 0.5 |
| Pan-Abeta | 4G8 | 800701 | Biolegend | 175 | Lu | 0.5 |
| PHFTau | PHF1_Tau | PHF1 | Peter Davis Lab | 176 | Yb | 0.25 |
| ***Primary Antibody** | | | | | | |
| TREM2 (biotinylated) | Poly | BAF1828 | R&D Systems | NA | NA | 2 |

#### Supplementary Table 3

| caseID | n_cells | AD | ApoE | gender | age | BraakScore | PlaqueTotal | TangleTotal |
|---|---|---|---|---|---|---|---|---|
| 11-107 | 1996 | yes | 4/4 | 1 | 71 | VI | 14 | 15 |
| 97-18 | 1107 | yes | 4/4 | 2 | 93 | VI | 14 | 14.5 |
| 07-21 | 464 | yes | 4/4 | 2 | 92 | VI | 14.5 | 15 |
| 11-78 | 1650 | yes | 3/3 | 1 | 82 | V | 14.5 | 15 |
| 10-46 | 1569 | yes | 3/3 | 1 | 86 | VI | 14 | 15 |
| 15-81 | 1329 | yes | 3/3 | 1 | 81 | VI | 14 | 14.5 |

**Supplementary Table 4**

| Antigen | Clone | Isotope | Element | Concentration (ug/ml) |
|---|---|---|---|---|
| p21-Waf1/Cip1 | 12D1 | 113 | In | 4 |
| PanAPOe | D7I9N | 141 | Pr | 2 |
| Abeta40 | BDI350 | 142 | Nd | 1 |
| p4EBP1 | 2855BF | 143 | Nd | 4 |
| p-PLCg2 | K86-689.37 | 144 | Nd | 2 |
| CD4 | RPA-T4 | 145 | Nd | 2 |
| Abeta42 | 12F4 | 146 | Nd | 1 |
| CD45 | D9M8I | 147 | Sm | 1 |
| CD83 | HB15e | 148 | Nd | 2 |
| p-CREB | polyclonal | 149 | Sm | 2 |
| CD11c | Bu15 | 150 | Nd | 2 |
| P2YR12 | polyclonal | 151 | Eu | 2 |
| Biotin | 1D4-C5 | 152 | Sm | 2 |
| TREM2-BIOTIN | polyclonal | 152 | Sm | 2 |
| CD14 | HCD14 | 153 | Eu | 2 |
| p-S6 | D57.2.2E | 154 | Sm | 0.5 |
| MerTK | Y323 | 155 | Gd | 1 |
| CD68 | D4B9C | 156 | Gd | 2 |
| GLUT5 | E-2 | 157 | Gd | 0.5 |
| gH2AX pSer139 | polyclonal | 158 | Gd | 1 |
| GLS | EP7212 | 160 | Gd | 1 |
| VDAC1 | 20B12AF2 | 161 | Dy | 1 |
| GPNMB | Poly | 162 | Dy | 1 |
| CD56 | NCAM16.2 | 163 | Dy | 1 |
| TMEM119 | CL8714 | 164 | Dy | 1 |
| p-p38 (T180/Y182) | 36/ p38 | 165 | Ho | 2 |
| IkBa | L35A5 | 166 | Er | 1 |
| ki67 | ki67 | 167 | Er | 4 |
| MRP14 | Poly | 168 | Er | 0.125 |
| Iba1 | EPR16588 | 169 | Tm | 1 |
| H3K9ac | C5B11 | 170 | Er | 1 |
| cPARP | Asp214 | 171 | Yb | 3 |
| H3K27me3 | C36B11 | 172 | Yb | 1 |
| CD44 | IM7 | 173 | Yb | 2 |
| HLADR | L243 | 174 | Yb | 1 |
| CD206 | 15-2 | 175 | Lu | 2 |

### Supplementary Table 5

**ezSegmenter segmentation parameters**

| mask_name | composite_add | composite_subtract | object_shape | sigma | thereshold | hole_size | min_size | max_size |
|---|---|---|---|---|---|---|---|---|
| amyloid_plaques | "Beta Amyloid 1-40", "Beta amyloid 1-42", | NA | blob | 1 | 85 | auto | 1000 | 100000 |
| tau | "PHF-tau" | NA | blob | 1 | 90 | auto | 200 | 100000 |
| microglia_arms | "P2RY12", "Iba1", "TMEM119" | NA | projection | 1 | 95 | auto | 300 | 100000 |
| astrocyte_arms | "S100B", "GFAP" | NA | projection | 0.7 | 85 | auto | 250 | 50000 |
| vessels | "CD31_CD105" | NA | projection | 2 | 98 | auto | 1000 | 100000 |
| neurons_soma | "Indium115" | "HistoneH3" | blob | 1 | 84 | auto | 1000 | 100000 |
| neurons_axons | "NEFH" | NA | projection | 2 | 95 | auto | 400 | 100000 |

**Mask merger parameters**

| first_merge | merge_masks_list | cell_mask_suffix | merge_operation | percent_overlap | output_mask_name |
|---|---|---|---|---|---|
| TRUE | ["neurons_axons"] | "neurons_soma" | combine | 1 | "neurons_combined" |
| TRUE | ["neurons_combined"] | "whole_cell" | combine | 90 | "neurons_combined_merged" |
| FALSE | ["microglia_arms", "astrocyte_arms"] | "whole_cell" | remove_duplicates | 20 | "microglia_arms_merged", "astrocyte_arms_merged" |

| **Parameter** | **Definition** |
|---|---|
| composite_add | Channels to add to composite channel. |
| composite_subtract | Channels to subtract from composite channel. |
| object_shape | General shape of the object, can be either `blob` or `projection` |
| sigma | The standard deviation for the Gaussian kernel to blur the image. Default is 1. |
| threshold | The per-fov-percentile threshold value (integer) for image thresholding. |
| hole_size | For any area smaller than `hole_size`, those holes are closed. Otherwise leave a 'auto' to determine the hole size based on image dimensions. |
| min_size | The minimum number of pixels required in a segmented object. |
| max_size | The maximum number of pixels permitted in a segmented object. |
| first_merge | Denotes if this is the first merge performed with a given merger base |
| merge_masks_list | List of object masks to merge to the base (cell) image. Merger occurs in order of appearance. |
| cell_mask_suffix | Suffix name of the cell mask files used as merger base. Usually `whole_cell` for Mesmer output. |
| merge_operation | `combine` integrates the base object into the final merged object. `remove_duplicates` identifies base objects that fulfill the merger criteria and removes the base object from the pool of base objects. |
| percent_overlap | Minimum percent threshold of a cell's area overlapping with an object that is required for a base cell mask to be merged into an object mask. |

**Round 1 merge**
Base input: mesmer nuclear output
Operation: `combine` with neurons
Output: (a) neurons_combined_merged.tiff (b) remaining_cells (unmerged mesmer outputs)

**Round 2 merge**
Base input: remaining_cells from round 1
Operation: `remove_duplicates` for microglia and astrocytes
Output: (a) microglia_arms_duplicates_removed.tiff (b) astrocyte_arms_duplicates_removed.tiff (c) remaining_cells (unmerged mesmer outputs)

### Supplementary Table 6

**Pathology and Neuronal Damage**

| *Feature* | *Description* | *Type* |
|---|---|---|
| tau_expansion_mean | Mean PHF-tau intensity in expansion zone (NFTs) | Pixel-level |
| amyloid_expansion_mean | Mean pan-amyloid intensity in expansion (plaques) | Pixel-level |
| tau_expansion_pos | Area positive for NFT (binary) | Object-level |
| amyloid_expansion_pos | Area positive for amyloid (binary) | Object-level |

**Metabolic Environment**

| *Feature* | *Description* | |
|---|---|---|
| ASCT2_expansion_mean | Glutamine transporter, metabolic activaiton zone | Pixel-level |
| GLS_expansion_mean | Glutaminase - supports metabolic rewiring | Pixel-level |
| ACC_expansion_mean | Fatty acid synthesis indicator | Pixel-level |
| LAL_expansion_mean | Lysosomal activity | Pixel-level |
| pS6_expansion_mean | mTOR pathway activation in environment | Pixel-level |

**Iron and Lysosome Biology**

| *Feature* | *Description* | |
|---|---|---|
| ferritin_expansion_mean | Iron buffering or storage in the neighborhood | Pixel-level |
| fe_expansion_mean | Free iron levels in the neighborhood | Pixel-level |

**Astrocyte and Glial Landscape**

| *Feature* | *Description* | |
|---|---|---|
| GFAP_expansion_mean | Reactive astrocyte load in expansion zone | Pixel-level |
| S100B_expansion_mean | Calcium-binding astrocyte marker | Pixel-level |
| CD44_expansion_mean | Upregulated in astrocyte-DAM crosstalk | Pixel-level |
| gliosis_index | Double postivity for GFAP and CD44 expansion expression | Object-level |

**Epigenetic and Stress Signals**

| *Feature* | *Description* | |
|---|---|---|
| pct_macroH2A1_pos | Presence of nucleated macroH2A1+ neighbor | Object-level |
| pct_gH2AX_pos | Presence of nucleated gH2A+ neighbor | Object-level |
| pct_H3K27me3_pos | Presence of nucleated H3K27me3+ neighbor | Object-level |
| pct_H3K9ac_pos | Presence of nucleated H3K9ac+ neighbor | Object-level |
| min_dist_to_gH2AX_pos | Distance to nearest gH2AX+ senescent cell | Object-level |

**Vascular/White Matter Proximity**

| *Feature* | *Description* | |
|---|---|---|
| CD31_CD105_expansion_mean | Endothelial markers | Pixel-level |
| CD56_expansion_mean | Seen on vasculature or neural adhesion sites | Pixel-level |
| MCNPase_expansion_pos | Area positive for MCNPase (binary) | Object-level |

**Neighborhood Composition**

| *Feature* | *Description* | |
|---|---|---|
| niche_entropy | Shannon entropy of neighbor types (log2) | Object-level |
| percent_tau_pos_neighbors | Percentage of neighbors that are tau+ | Object-level |
| percent_amyloid_pos_neighbors | Percentage of neighbors that are amyloid+ | Object-level |
| min_dist_to_tau | Distance to nearest segmented tau+ object (if available, including neuror | Object-level |
| min_dist_to_amyloid | Distance to nearest segmented amyloid object (if available) | Object-level |
| min_dist_to_vessel | Distance to nearest vessel object (if available) | Object-level |

#### APOE4-associated signature

| | | | | | | | | | | |
|---|---|---|---|---|---|---|---|---|---|---|
| FTH1 | FTL | EEF1A1 | MT-ATP6 | RPL19 | MT-CO2 | PTMA | CD74 | CD81 | C1QA | MT-CO3 |
| C1QB | RPS2 | C1QC | ACTB | HLA-DRA | APOE | TPT1 | MT-CYB | RPL10 | MT-ND3 | CST3 |
| RPL15 | RPS3A | RPL13 | MT-ND2 | RPS18 | CD14 | RPL13A | B2M | MRC1 | RBMS3 | HLA-DRB1 |
| RPL30 | TMBIM6 | MT-ND4 | RPL35 | MT-ND1 | RPS19 | MT-CO1 | CD63 | RPL36A | RPL41 | RPL29 |
| TUBA1B | HLA-A | RPL18A | RPL28 | RPS27A | RPS14 | RPLP1 | RPS15 | RPL18 | RPL17 | H3F3B |
| VSIG4 | RPL7A | HLA-C | NPC2 | RPL32 | HLA-B | FAU | RPL8 | SERF2 | RPS3 | HLA-E |
| RPLP2 | RPL24 | PSAP | RPL9 | RPL6 | TMSB4X | RPL11 | RPL3 | LINC00486 | HLA-DPA1 | GRID2 |
| RPS8 | SLC9A9 | LINC01684 | RPS6 | RPL5 | RPS20 | USP53 | TRPS1 | RPS24 | MERTK | |
